# Imputation of low-coverage sequencing data from 150,119 UK Biobank genomes

**DOI:** 10.1101/2022.11.28.518213

**Authors:** Simone Rubinacci, Robin Hofmeister, Bárbara Sousa da Mota, Olivier Delaneau

## Abstract

Recent work highlights the advantages of low-coverage whole genome sequencing (lcWGS), followed by genotype imputation, as a cost-effective genotyping technology for statistical and population genetics. The release of whole genome sequencing data for 150,119 UK Biobank (UKB) samples represents an unprecedented opportunity to impute lcWGS with high accuracy. However, despite recent progress^1,2^, current methods struggle to cope with the growing numbers of samples and markers in modern reference panels, resulting in unsustainable computational costs. For instance, the imputation cost for a single genome is 1.11£ using GLIMPSE v1.1.1 (GLIMPSE1) on the UKB research analysis platform (RAP) and rises to 242.8£ using QUILT v1.0.4. To overcome this computational burden, we introduce GLIMPSE v2.0.0 (GLIMPSE2), a major improvement of GLIMPSE, that scales sublinearly in both the number of samples and markers. GLIMPSE2 imputes a low-coverage genome from the UKB reference panel for only 0.08£ in compute cost while retaining high accuracy for both ancient and modern genomes, particularly at rare variants (MAF < 0.1%) and for very low-coverage samples (0.1x-0.5x).

## Main

To demonstrate the benefits of using sequenced biobanks for lcWGS imputation, we phased the recent release of the UK Biobank (UKB) WGS data^3,4^ using SHAPEIT5^5^ and created a UKB reference panel of 280,238 haplotypes and 582,534,516 markers (**Supplementary Note S1**). We used the UKB panel to impute lcWGS samples with GLIMPSE2 and other recently released imputation methods: GLIMPSE1^1^ and QUILT v1.0.4^2^. Compared to other reference panels, the UKB leads to considerable accuracy improvements for British samples across all tested depths of coverage. Furthermore, GLIMPSE2 outperforms GLIMPSE1, particularly at rare variants (MAF<0.1%) and for very low-coverage (0.1-0.5x) and matches QUILT v1.0.4 accuracy, designed to condition on the full set of reference haplotypes (**Figure 1a, Supplementary Note S2**). To consider non-British populations, we imputed 276 lcWGS samples from the Simons Genome Diversity Project and we show that the UKB panel drastically improves imputation accuracy of European samples, in particular of Northern Europe origin (**Supplementary Note S3**). Additionally, we imputed three ancient Europeans and a Yamnaya sample for which high-coverage data (>18x) is available, and find similar improvements (**Supplementary Note S4**), showing that some ancient populations, such as Viking, Western Hunter-Gatherer and Yamnaya could be well imputed from the UKB reference panel.

**Figure 1:**
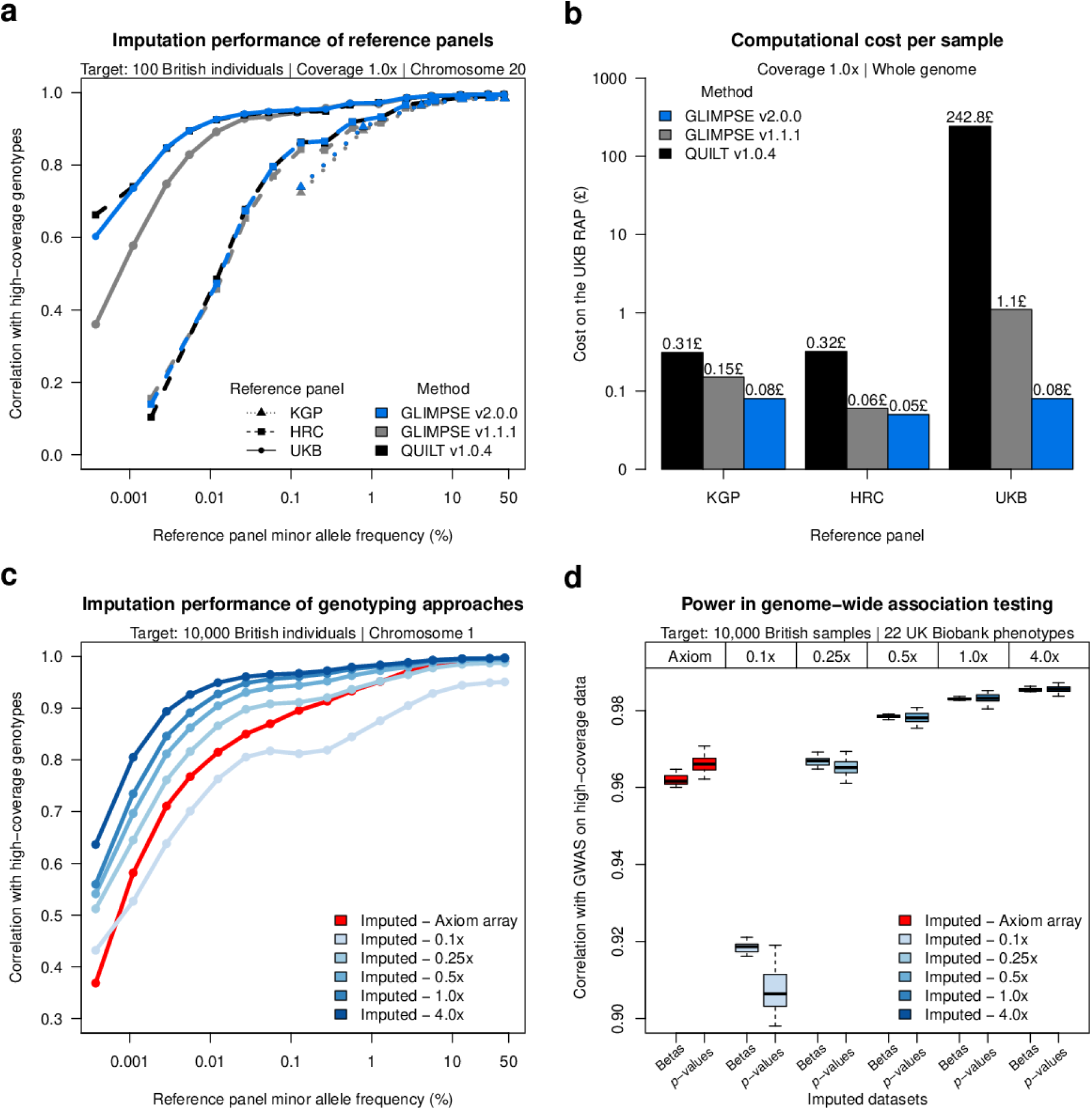
Accuracy, running time and power of low-coverage imputation using the UK Biobank WGS data. (**a**-**b**) Imputation performance of different imputation methods: QUILT v1.0.4 (black), GLIMPSE1 (grey) and GLIMPSE2 (blue); across different reference panels (KGP, HRC and UKB) for 100 UKB British samples at 1.0x coverage. (**a**) Accuracy on chromosome 20 (Pearson *r*^2^, y-axis), of imputation methods and reference panels: KGP (triangle dotted line), HRC (squared dashed line) and UKB (full line). Accuracy in plotted against minor allele frequency of the appropriate reference panel (x-axis, log-scale). (**b**) Cost per sample on the RAP for whole-genome imputation (y-axis, log scale) across different reference panels (x-axis). (**c**-**d**) Performance of imputed data using the UKB reference panel across coverages (0.1-4.0x, different shades of blue, GLIMPSE2 imputation), and Axiom array data (red). (**c**) Accuracy on chromosome 1 of 10,000 UKB British samples (Pearson *r*^2^, y-axis) against minor allele frequency of the appropriate reference panel (x-axis, log-scale). (**d**) Power in GWAS in association testing of 10,000 UKB British samples compared to high-coverage data. Correlation of betas and p-values (y-axis) of different imputed datasets (x-axis) across 22 UKB phenotypes.

The imputation of a single lcWGS genome using the UKB reference panel is expensive using existing methods. On the RAP, the cost is 1.11£ and 242.80£ for GLIMPSE1 and QUILT v1.0.4, respectively. In contrast, the same task performed with GLIMPSE2 only costs 0.08£, due to major algorithmic improvements that drastically reduce the imputation time for rare variants (**Fig 1b, Supplementary Note S2**). We confirm this trend for up to 2 million reference haplotypes, using simulated data. This keeps lcWGS close to SNP arrays in terms of computational costs^6^ (**Supplementary Note S3**) while having the major advantage of providing better genotype calls. Indeed, we find that imputation of 0.5x data yields to more accurate results than the UKB Axiom array, specifically designed for the British population, with a notable difference at rare variants (for 0.5x coverage, accuracy improvement of *r*^2^ > 0.1 for variants with a MAF < 0.01%, **Figure 1c**).

To assess the impact of these improvements on Genome-Wide Association Studies (GWAS), we imputed 10,000 UKB samples that we used to test 22 quantitative traits for association, comparing the respective abilities of lcWGS and SNP array data to recover the signals found with high-coverage sequencing data. We find that 0.25x leads to p-values and effect size estimates as accurate as those obtained from Axiom array data (**Figure 1d**) while delimiting regions of association with matching sensitivity and specificity (**Supplementary Note S6**). We also look at rare loss-of-function, missense and synonymous variants^7^, and show that 1.0x significantly outperforms the Axiom array in burden-test analysis (**Supplementary Note S7**). Altogether, this shows that lcWGS constitutes a powerful alternative to SNP array for downstream GWAS and rare variant analysis.

In this work, we introduce several major improvements for GLIMPSE that solve the computational problem of imputing lcWGS data from the 150,119 WGS samples in the UK Biobank. We demonstrate that this new reference panel leads to striking accuracy improvements across all allele frequencies, sample ancestries, and depths of coverages. Our study further confirms the advantage of lcWGS over SNP arrays for GWAS, by showing that using imputed data with coverage as low as 0.5x is enough to outperform a SNP array specifically designed for the target population, particularly at rare variants. Our work can be applied to other sequenced and diverse biobanks, such as Trans-Omics for Precision Medicine^8^, gnomAD^9^ or AllofUs^10^, thereby facilitating lcWGS imputation of non-European individuals. We believe that the difference between low-coverage and high-coverage WGS will become increasingly smaller as large reference panels will keep collecting more human haplotype diversity.

## Code availability

GLIMPSE2 source code is available with MIT licence from https://github.com/odelaneau/GLIMPSE and https://odelaneau.github.io/GLIMPSE/. Pre-compiled binaries and docker images are available at https://odelaneau.github.io/GLIMPSE/release. Scripts to produce all figures of the paper are available on Github.

## Data availability

The 1000 Genomes Project phase 3 dataset sequenced at high coverage by the New York Genome Center is available on the European Nucleotide Archive under accession no. PRJEB31736, the International Genome Sample Resource (IGSR) data portal and the University of Michigan school of public health ftp site (URL: ftp://share.sph.umich.edu/1000g-high-coverage/freeze9/phased/). The publicly available subset of the HRC dataset is available from the European Genome-phenome Archive at the European Bioinformatics Institute (EBI) under accession no. EGAS00001001710. The publicly available Simons Genome Diversity project is available on the IGSR data portal and Cancer Genomics Cloud, powered by Seven Bridges. The UK Biobank genetic and phenotypic data are available under restricted access. Access can be obtained by application via the UK Biobank Access Management System (URL: https://www.ukbiobank.ac.uk/).

## Acknowledgements

This work was funded by a Swiss National Science Foundation project grant 373 (PP00P3_176977) and conducted under UK Biobank project 66995. The New York Genome Center 1000 Genomes data were generated at the New York Genome Center with funds provided by a National Human Genome Research Institute grant no. 3UM1HG008901–03S1.

## Contributions

S.R. and O.D. designed the study. S.R. and O.D. developed the algorithms and wrote the software. R.J.H. performed the GWAS experiments. S.R. and B.S.M performed imputation of ancient samples. B.S.M. provided interpretation regarding imputed ancient samples. S.R. performed the remaining experiments. O.D. supervised the project. All authors reviewed the final manuscript.

## Corresponding author

Correspondence to Olivier Delaneau.

