## Supplementary Notes for "Imputation of low-coverage sequencing data from 150,119 UK Biobank genomes"

### Supplementary Information for “Imputation of low-coverage sequencing data from 150,119 UK Biobank genomes”


**Supplementary Note 1. Method and data description** 2
- S1.1 Data 2
- S1.2 GLIMPSE version 2.0.0 4
- S1.3 Benchmark description 10

**Supplementary Note 2. Performance of low-coverage imputation methods** 14
- S2.1 Introduction 14
- S2.2 Methods 14
- S2.3 Results 15

**Supplementary Note 3. Imputation performance of low-coverage and SNP arrays** 17
- S3.1 Introduction 17
- S3.2 Methods 17
- S3.3 Results 17

**Supplementary Note 4. Imputation of Simons Genome Diversity Project dataset** 19
- S4.1 Introduction 19
- S4.2 Methods 19
- S4.3 Results 19

**Supplementary Note 5. Imputation of ancient European populations** 21
- S5.1 Introduction 21
- S5.2 Methods 21
- S5.3 Results 22

**Supplementary Note 6. Common-variant association analyses** 23
- S6.1 Introduction 23
- S6.2 Methods 23
- S6.3 Results 24

**Supplementary Note 7. Rare-variant association analyses** 25
- S7.1 Introduction 25
- S7.2 Methods 25
- S7.3 Results 26

**References** 27

**Supplementary Figures** 31

**Supplementary Tables** 46

### **Supplementary Note 1.** Method and data description

### S1.1 Data

#### S1.1.1 UK Biobank

The UK Biobank (UKB) is a resource containing phenotypic and genotypic data across ~500,000 individuals from the UK. The first release of the data was generated using SNP arrays for all participants[1], but genotypes and phenotypes are constantly updated and evolving. The UKB is currently sequencing the genomes of all the participants to an average coverage of 32.5× (at least 23.5× per individual). A first data release of the whole genome sequencing (WGS) data for 150,199 individuals has been made available[2], and consists of a large set of sequence variants, SNPs, indels, microsatellites and structural variants. Sequence reads have been mapped to the human reference genome GRCh38. Variant discovery process and genotype calling has been jointly performed across individuals.

In our analysis, we use WGS data jointly called with GraphTyper[3] and made available as pVCF files on the UK Biobank RAP. We performed quality control on the WGS data as follows. First, we decomposed multi-allelic variants into bi-allelic variants using BCFtools[4], then we filtered out SNPs and indels for (i) Hardy-Weinberg p-value < $10^{-30}$, (ii) more than 10% of the individuals having no data (GQ score=0; i.e. missing data), (iii) excess of heterozygosity, quantified to be less than 0.5 or greater than 1.5, (iv) alternative alleles with AAscore < 0.5, and (v) variant sites with the tag FILTER different from PASS. This resulted in a total of 603,925,301 variant sites, including 20,662,402 common variant sites (MAF ≥ 0.1%) and 583,262,899 rare variant sites (MAF < 0.1%), across a total of 150,119 individuals. We then performed haplotype phasing on all autosomes using SHAPEIT5[5]. We extracted 10,000 samples with British ancestry that are unrelated to any other sample in the dataset and for which we had SNP array data available for the purpose of imputation experiments. These constitute the target samples, for which we downsampled sequencing data at multiple low coverages. The remaining 140,119 WGS samples constitute a reference panel variable at 582,534,516 sites (**Supplementary Table 1**).

#### S1.1.2 1000 Genomes Project

The 1000 Genomes Project[6] (KGP) was designed to provide a comprehensive catalogue of human genetic variation across different ancestries. The phase 3 of the project contains sequenced data of

2,504 individuals sampled from 26 different populations that can be divided into five continental groups: 661 samples with African ancestry (AFR), 347 samples with American ancestry (AMR), 503 samples with European ancestry (EUR), 504 samples with East Asian ancestry (EAS) and 489 samples with South Asian ancestry (SAS). Recently, the New York Genome Center (NYGC), funded by National Human Genome Research Institute, has resequenced all 2,504 samples at 30x depth of coverage, including data of 602 closely related individuals[7]. Sequencing was performed using Illumina NovaSeq 6000 with 2×150bp reads that were aligned to GRCh38. The data is shared via the International Genome Sample Resource (IGSR)[8].

In this work, we used a version of the KGP phase 3 project from which variant discovery and haplotype phasing has been performed as part of a larger collection of 143,415 participants across 37 TOPMed studies, and referred to as "data freeze 9". The data, for which quality control and haplotype phasing has been performed by the data providers, is publicly available (URL). Information about the number of variants and samples used are shown in **Supplementary Table 1**.

##### S1.1.3 Haplotype reference consortium

The Haplotype Reference Consortium (HRC)[9] is a phased reference panel obtained from sequence data of 32,470 individuals from multiple cohorts with relatively low coverage WGS (from 4x to 8x coverage). The panel contains a set of 39,235,157 variants (bi-allelic SNPs) with minor allele counts (MAC) ≥ 5, collected from 20 different studies (mostly European ancestry). The publicly available version of the data contains 22,691 individuals (chromosome 1) and 27,165 individuals (chromosomes 2-22, X), with phased SNP genotypes aligned to GRCh37.

This version of the HRC data was downloaded from the European Genome-Phenome Archive at the European Bioinformatics Institute (accession EGAS00001001710). We used the Picard toolkit (URL) to liftover the data to the human reference genome GRCh38. We discarded data for strand flips, obtaining 99.8% of the original variants after the liftover. Information about the number of variants and samples used are shown in **Supplementary Table 1**.

##### S1.1.4 Simons Genome Diversity Project

The Simons Genome Diversity Project[10] (SGDP) is a dataset containing 300 samples from more than one hundred diverse human populations. Of these 300 samples, 279 complete genomes have been made publicly available on the Cancer Genomics Cloud, powered by Seven Bridges (URL). Additionally, the EBI made available a version of the data for 276 individuals out of the 279, aligned to GRCh38 (URL).

The 276 samples come from 129 different populations across the world: 44 Africans, 22 Native Americans, 27 Central Asians or Siberians, 47 East Asians, 25 Oceanians, 36 South Asians and 75 West Eurasians. All genomes in the dataset were sequenced to at least 30x coverage using Illumina technology. The data has been mapped and genotyped using a customised procedure that was optimised for population genetic analysis.

In order to generate validation data for our imputation experiments on chromosome 20, we used high confidence genotype calls at variable sites produced by the authors in VCF format, and filtered genotypes with FORMAT/FL value greater or equal to zero, following the recommendations of the authors. We then lifted over to GRCh38 and intersected the variants at known sites in all the UKB, KGP and Gnomad v3.1.2 datasets, obtaining a total of 510,809 variants on chromosome 20. For the imputation experiments we used downsampled data from the GRCh38 alignment files available on the EBI website.

### S1.2 GLIMPSE version 2.0.0

#### S1.2.1 Background

GLIMPSE version 2.0.0 (GLIMPSE2) is a method for haplotype phasing and genotype imputation for low-coverage whole genome sequencing (lcWGS) samples. The input of the algorithm is primarily made of two components: a reference panel $H$, defined on $N$ haplotypes and $M$ markers, and a matrix of genotype likelihoods for the low-coverage samples to impute, called targets or target samples in the following. Genotype likelihoods are defined as the probability of the observed sequencing reads given the unknown genotype, here assumed to be biallelic. In the context of imputation, genotype likelihoods are restricted to the set of $M$ markers in the reference panel.

#### S1.2.2 Model description

GLIMPSE2 is a method that performs haploid imputation and diploid phasing in a Gibbs sampler scheme, starting from a matrix of genotype likelihoods for the target samples and a reference panel of haplotypes. As the haplotype-level data is unknown at the beginning of the algorithm, a first initialising iteration is run to provide a first haplotype estimate. GLIMPSE2 selects an initial set of haplotypes (default K_init=1000), from the reference panel, and runs two consecutive steps of haploid imputation, one for each of the two target haplotypes. The two haplotypes are then sampled from the imputation posteriors. The haploid imputation step relies on a modified version of Li and Stephens hidden Markov model (HMM)[11] that uses haploid likelihoods as the emission layer. GLIMPSE2 derives haploid likelihoods for one of the two target haplotypes, from the original genotype likelihoods, conditional to the current estimate of the other haplotype. At the end of the initialization step, a diplotype is assigned to each target sample. GLIMPSE2 then runs a set of burn-in (5 by default) and main Gibbs iterations (15 by default) to refine the imputation and phasing of each target sample. The algorithm averages imputation posteriors across all main iterations, to integrate over phasing uncertainty. More information regarding the Gibbs sampling scheme can be found in the original GLIMPSE paper[12].

GLIMPSE2 is designed to perform imputation based only on the reference panel, and optimises this task with seven novel key features:

1. A sparse memory representation of the reference panel that stores efficiently the large number of rare variants it contains (**Section S1.2.2.1**),
2. An efficient implementation of the hidden Markov model (HMM) that speeds up probability computations by leveraging the sparsity of the reference panel (**Section S1.2.2.2**),
3. A new data structure based on the Positional Burrows Wheeler Transform (PBWT) that speeds up haplotype matching by leveraging the sparsity of the reference panel (**Section S1.2.2.3**),
4. A sparse file format for the reference panel, containing also the pre-computed PBWT data structures, that allows fast loading times (**Section S1.2.2.4**),
5. A genotype caller that internalises the pile-up of the sequencing data and the computations of genotype likelihoods (**Section S1.2.2.5**),
6. A model extension to impute small indels and low-quality bi-allelic variants separately from SNPs (**Section S1.2.2.6**),
7. An optimised iteration scheme that integrates an initialisation step based on rare variant sharing (**Section S1.2.2.7**).

###### S1.2.2.1 Sparse reference panel representation

GLIMPSE2 is designed for large reference panels containing hundreds of millions of genetic variants, many of which are rare. For this reason, we developed a sparse matrix representation of the reference haplotypes to efficiently store large amounts of data in memory and speed up computations. The memory representation of GLIMPSE2 considers each marker independently, and it represents reference haplotypes in one of the following two ways: if the minor allele is rare (MAF < 0.001 by default), the indices of the haplotypes that carry the minor allele are stored, otherwise the full sequence of alleles is stored using one bit per allele. This representation allows for a large reduction in memory usage and a fast identification of the haplotypes carrying a rare variant. Additionally, by transposing this data structure, we can quickly access all rare variants of each reference sample. In practice, this representation allows the model to reduce memory requirements, speed up calculations and scale to larger imputation windows.

**S1.2.2.2 Sparse hidden Markov model computations**

GLIMPSE2 uses the Li and Stephens HMM[11] to perform imputation and phasing. Here we briefly introduce the standard approach to perform imputation on low-coverage WGS data, and later introduce a few computational speedups obtained by the reference panel representation.

To calculate the probability of a target haplotype $t$, observed at $M$ markers, the model works on a subset $X$ of reference haplotypes, selected using the PBWT, defining the hidden states of the HMM. Let $X$ be defined as the subset of haplotypes $h_i$, $i < |X|$, over the $M$ markers, $h_i = \{h_{i,0}, h_{i,1}, \dots, h_{i,M-1}\}$, where $h_{i,m} \in \{0, 1\}$. We use the forward-backward algorithm[11,13,14] to compute posterior probabilities for each of the states of the HMM. This results in a probability of each hidden state given the target haplotype, being:

$$P(X_m \mid t) \propto \alpha_m(h_i)\beta_m(h_i) .$$

Posterior allele probabilities are computed by summing state probabilities for reference haplotypes that carry the same allele. Genotype probabilities are obtained by combining posterior probabilities of the two haplotypes of the same sample.

We designed the GLIMPSE2 algorithm to remove the linear component at the marker level during the HMM calculations by leveraging the sparsity of the reference panel at rare variants. Indeed, as GLIMPSE2 selects only $K$ (default $K = 2000$) haplotype samples with the sparse PBWT selection (**Section S1.2.2.3**), most of the rare variants present in the reference panel are not variable in the custom reference panel. In the forward-backward algorithm these monomorphic variants do not contribute to the overall state probability. Therefore, we compute the forward-backward probabilities only at sites that are polymorphic in the custom reference panel, by adjusting the transition probability to consider the distance in Morgan between two consecutive polymorphic sites. Posterior probabilities of variants that are monomorphic in the composite reference panel can be quickly computed by using the appropriate emission probability. We efficiently test if the custom reference panel is monomorphic using the transpose of the reference panel data structure at rare variants.

Additionally, we take advantage of low-level programming language (AVX2 intrinsics) to optimise the Li and Stephens forward-backward computations at the hardware level. The forward computations can be implemented with a fused multiply add (FMA) operation for the transition and a SIMD multiplication based on the mask as the emission layer. A similar implementation can be used for the backward calculations. By working on blocks of 8 floats, we obtain a substantial speedup of the forward-backward algorithm, allowing GLIMPSE2 to use twice the number of states and larger imputation windows than the previous version of GLIMPSE.

**S1.2.2.3 State selection using the sparse PBWT**

The GLIMPSE1 model reduces the state space for imputation and phasing by creating the positional Burrows Wheeler transform (PBWT)[15] of the reference haplotypes and quickly extracting haplotypes sharing long matches with the target haplotypes. Haplotypes that share long identity-by-state segments cluster together in the data structure. Haplotype selection exploits this fact to efficiently select sets of highly informative haplotypes for imputation and phasing. GLIMPSE2 introduces a new data structure for state selection, called sparse PBWT, that exploits the sparse reference panel representation to speed up PBWT computations in the presence of large amounts of rare variants.

The PBWT is a transformation applied to each column of a binary matrix representing the reference panel $H$. Let $H$ be defined on a set of N haplotypes $h_n$, $n < N$, over $M$ markers, $h_n = \{h_{n,0}, h_{n,1}, \dots, h_{n,M-1}\}$, where m < M, and $h_{n,m} \in \{0, 1\}$. As $H$ is a N × M binary matrix we refer to $H$ using the usual matrix notation. The PBWT of $H$, indicated as $Y$, is another $N \times M$ binary matrix, where the $m$-th column of Y is an invertible transformation of the $m$-th column of $H$ representing the haplotypes in the order of the reversed prefixes. In its basic form, $Y$ is complemented by another matrix $A$, where every column $A_{:,m}$, called the positional prefix array at marker $m$, is a permutation of $\{0, N-1\}$ which defines the reverse prefix order of the haplotypes in $H$ up to marker $m-1$. Let $A_{:,0}$ be the identity permutation. Efficient algorithms to compute $Y$ and $A$ are provided in the original PBWT publication[15].

Here we propose an algorithm to compute the positional prefix array and haplotype matching that is designed for large sequencing cohorts. The key idea is that, by using the sparse representation of the reference panel, we can proceed differently for common and rare variants, computing the standard PBWT update at common variants and work on smaller PBWTs at rare variants. First, we note that between two adjacent common variants most of the haplotypes do not contain the minor allele in the region, and therefore most of the haplotypes would form a single invariable block of major alleles that preserves their relative haplotype order. A smaller PBWT is constructed only on haplotypes that have at least one minor allele between two adjacent common variants. The positional prefix array of the small PBWT at the end of the rare-variant interval is enough to reconstruct the full prefix array at the next common variant. The prefix array of the small PBWT is simply concatenated with the positional prefix array of other haplotypes that are not changing in the interval. A schematic illustration of the sparse PBWT is shown in **Supplementary Figure 1**.

More formally, let us consider the reference panel $H$ bounded between consecutive common variants $P$ and $Q$. Let $A_{:,P+1}$ be the positional prefix array at marker $P$, and $A_{:,Q}$ be the positional prefix array at marker $Q-1$. Let us consider $S_{P,Q}$ to be the set haplotypes containing at least one minor allele in the region between markers $P$ and $Q$ (excluded), obtained simply taking the union of the lists provided by the sparse reference panel representation, and define its size as $L = |S_{P,Q}| \leq N$. We first note that all haplotypes not in the set $S_{P,Q}$ are not variable in the region, and therefore they will maintain the same relative prefix order of $A_{:,P}$ and form a block at the beginning of $A_{:,Q-1}$ of size $N-L$. Second, we complement $A_{:,Q-1}$ with the haplotypes in $S_{P,Q}$, by considering a PBWT computed only for haplotypes in $S_{P,Q}$ In practice, we consider the sparse positional prefix array $A'_{:,P+1}$ at marker $P$ (of size $L$) by taking all haplotypes in $S_{P,Q}$ in the order of $A_{:,P+1}$. Starting from $A'_{:,P+1}$ construct the sparse positional prefix array at all markers between $P$ and $Q$ (excluded): $A'_{:,P+1}, A'_{:,P+2}, A'_{:,P+3}, \dots, A'_{:,Q-1}$ using the standard algorithms. At the end of this process, $A'_{:,Q-1}$ can be concatenated to the invariant part of $A_{:,Q-1}$ from position $(N-L)$. This is equivalent to a standard PBWT update between $P$ and $Q$ (excluded). We can therefore proceed to compute the standard PBWT update at marker $Q$.

The algorithm builds on the idea that the number of common variants (e.g., with MAF > 0.1%) remains approximately constant as the number of reference samples increases, while the number of rare variants keeps growing. Haplotype selection is performed using algorithm 5 in the original PBWT algorithm[15], querying prefix arrays at common variants (at 0.1 cM intervals by default). The selection is complemented with variant sharing at rare variants, as rare variant sharing is likely to arise from a recent common ancestor[16].

**S1.2.2.4 File format**

We use a binary file format to store the sparse representation of the reference panel (**Section S1.2.2.1**) on the disk. The file format translates directly into the memory data structures used by GLIMPSE2 and does not require any compression algorithms such as GZIP. Additionally, GLIMPSE2 skips redundant computational steps by storing the sparse PBWT data structure in the same file format, together with the recombination map. As a result, all the data related to the reference panel can be quickly loaded in memory, in much faster running times than standard file formats, such as VCF and BCF.

**S1.2.2.5 Internal genotype likelihood computation**

GLIMPSE2 can call genotype likelihoods from raw sequencing read data in BAM/CRAM file format, using the GATK (dragon) model that is defined as:

$$P(G = \{a_1, a_2\}) = \prod_{k=1}^{M} \quad P(b_k | G = \{a_1, a_2\}) = \prod_{k=1}^{M} \quad \left(\frac{1}{2} P(a_1) + \frac{1}{2} P(a_2)\right)$$

$$P(a) = \{\frac{e}{3} \;\; if\ b \neq a; \quad 1 - e \quad otherwise;$$

where M is the sequencing depth at the site, $b_k$ is the observed base in read $k$, $e$ is the probability of error calculated from the phred scaled quality score of the base ($e = \ 10^{-\frac{q}{10}}$).

As this model might not perform well for higher sequencing depths and complex genetic variation, GLIMPSE2 also offers the possibility to read pre-computed genotype likelihoods in VCF/BCF format in order to take advantage of more sophisticated genotype callers.

**S1.2.2.6 Imputation at small indels and low-quality variants**

Imputation of small indels and low-quality variants can be challenging for imputation methods as errors made at these markers can propagate to neighbouring makers and lower imputation quality. To prevent this, GLIMPSE2 uses an approach based on haplotype scaffolding[17]. First, indels and any other low-quality variants are flagged, leaving bi-allelic SNPs as high-quality variants. Then, the HMM forward-backward computations are performed only on high quality variants, ignoring low-quality variants and preventing errors at these variants to propagate to neighbouring ones. This builds a haplotype scaffold at bi-allelic SNPs onto which low-quality variants are phased and imputed in turn, in a similar way to what is done in the context of SNP-array data imputation[13].

When imputation is performed into the scaffold, genotype likelihoods are used as an emission of the model. The internalised genotype likelihoods model of GLIMPSE2 does not offer a well-calibrated model to confidently call genotype likelihoods at INDEL sites. For this reason, the default behaviour of GLIMPSE2 is to assume missing genotype likelihoods at INDEL sites.

**S1.2.2.6 Iteration scheme and initialisation using rare-variant sharing**

At the start of a run, GLIMPSE1 selects a fixed number of random reference haplotypes, used for a first imputation and phasing pass. However, as reference panels grow, using random haplotypes becomes sub-optimal. Based on previous work on phasing[16,18], we propose a new initialisation strategy that selects reference haplotypes sharing rare alleles with the target sample.

As we work with genotype likelihoods, we first perform genotype calling using as a prior the minor allele frequency of the variant sites in the reference panel[19]. This gives us fixed genotype calls that we use for haplotype selection in the reference panel: we focus on genotypes carrying at least a copy of the minor allele and search for candidate haplotypes in the reference panel carrying the same minor allele. This is performed efficiently using the sparse reference panel data structure. In practice, we fill the list of candidate haplotypes by considering the rarest variant sites first, until 800 initialising haplotypes have been selected. Then, we complete the list with an additional 200 haplotypes that we randomly pick in the reference panel. The resulting 1,000 haplotypes are finally used to carry out the first imputation and phasing pass of the data and obtain a first haplotype estimate for all target samples.

Finally, GLIMPSE2 treats burn-in iterations differently from main iterations. During the burn-in step, GLIMPSE2 runs two consecutive imputation steps, the first with a smaller number of states ($K/5$), and the second with the full $K$, selected from the sparse PBWT. This allows the method to have a better phasing at common variants prior to the full iteration. During the main iterations, genotype posteriors are stored. The final genotype posteriors are a result of the average across all main iterations.

### S1.3 Benchmark description

#### S1.3.1 Measurements of imputation accuracy

We measured imputation performance as the squared Pearson correlation between the validation (high coverage) and the imputed dosages. We computed imputation performance as a function of minor allele count, that we computed from the high-coverage reference panels. For the UK reference panel, we classified all variants into the following bins of minor allele count:

[1,5], [5,10], [10,20], [20,50], [50,100], [100,200], [200,500], [500,1000] [1000,2000], [2000,5000], [5000,10000], [10000,20000], [20000,50000], [50000,100000], [100000,140119].

For smaller reference panels, only an appropriate subset of the bins was considered. To get reliable measures of imputation performance, we pooled together all validation and imputed dosages belonging to the same frequency bin and computed a single squared Pearson correlation value per bin. Statistics summarising the number of variants falling in each allele count bin are provided in (**Supplementary Table 2-4**). We used the GLIMPSE2_concordance tool to measure the squared Pearson correlation by streaming the imputed and validation data to maintain low memory requirement. We extended the GLIMPSE2_concordance too to output an $r^2$ per allele frequency bin depending on the variant sites, getting an $r^2$ value per variant type (SNPs, INDELs and SNPs/INDELs combined).

We also used the GLIMPSE2_concordance tool to measure the accuracy at the genotype level, computing the genotype discordance between imputed and validation genotypes at the sample level or MAF bins (homozygous major; heterozygous; homozygous minor).

Finally, we also evaluated the non-reference discordance rate. The non-reference discordance rate is calculated as $nrd = \frac{(100 \times (e_{rr}+e_{ra}+e_{aa}))}{(e_{rr}+e_{ra}+e_{aa}+ m_{ra}+m_{aa})}$, where $e_{rr}$, $e_{ra}$ and $e_{aa}$ are the counts of the mismatches for the homozygous reference, heterozygous and homozygous alternative genotypes, while $m_{ra}$ and $m_{aa}$ are the counts of the matches at the heterozygous and homozygous alternative genotypes. We define the non-reference concordance rate as: $nrc = 100 - nrd$.

#### S1.3.2 Low-coverage imputation methods and parameters

For the benchmarks, we focused on three imputation methods: GLIMPSE v1.1.1[12] (release May 2021; here called GLIMPSE1), QUILT v1.0.4[20] (release November 2022) and GLIMPSE v2.0.0 (here called GLIMPSE2). We ran imputation analysis using the same window size of 2Mb for GLIMPSE1 and QUILT, as for a total of 32 chunks on chromosome 20, with a buffer region of 250kb. For GLIMPSE2, we used chunks of 4Mb as the method is designed to scale to larger imputation windows. We generated chunks using the GLIMPSE2_chunk tool. All methods were run using 4 threads, except otherwise stated. We used default parameters for every method. Programs were run in a docker container on the RAP.

**S1.3.2.1 GLIMPSE v2.0.0 [GLIMPSE2]**

We first split the reference panel into 4Mb chunks and created a binary representation of the file format, together with the recombination map and sparse PBWT using the split_reference tool. We used GLIMPSE2 to perform imputation with default parameters, using a single job for each chunk and providing a list of BAM files directly to the software as an input. After the imputation step was completed, we ligated the chunks using the ligate tool with default parameters, to get a single file for each chromosome.

**S1.3.2.1 GLIMPSE v1.1.1 [GLIMPSE1]**

We used GLIMPSE v1.1.1 with default parameters, using a job for each chunk. After the imputation step was completed, we ligated the chunks using the GLIMPSE1_ligate tool with default parameters, to get a single file for each chromosome. We note that the ligate tool in v1.1.1 differs from the ligate tool in v2.0.0 as GLIMPSE2 does not output the FORMAT/HS field.

**S1.3.2.1 QUILT v1.0.4 [QUILT]**

We converted the reference panel into the IMPUTE format using BCFtools. Then, we created the binary representation of the reference panel for QUILT using the QUILT_prepare_reference.R script with default options and the genetic map converted in the QUILT format. Finally, we run an imputation using the QUILT.R script, specifying the list of BAM files, the start/end region and buffer length, four cores and otherwise default parameters. We ligated imputed chunks using BCFtools as recommended by the authors.

##### S1.3.3 Running methods on the RAP

To run a computational job on the RAP, the first step is to choose a Virtual Machine (VM). VMs differ in the amount of storage (ssd 1-3), RAM (mem 1-3) and vCPUs (x 2-128) they contain and their respective price is available on the DNA Nexus website (URL). For imputation from small reference panels (KGP and HRC), all methods run on the smallest VM with four vCPUs. For the UKB reference panel, methods required different hardware settings and we matched as much as possible the VMs to the minimal hardware required by the respective methods (**Supplementary Table 5-7**).

The RAP is based on Amazon Web Services and offers a choice of two types of allocated instances for computations, “spot” (lower cost) and “on-demand” (higher cost), depending on the priority given to the jobs. There are three priority levels for jobs on the RAP:

1. **low**: tries to run the job on “spot” instances but the computation can be interrupted by the scheduler.
2. **normal**: jobs will start executing on “spot” VMs, but this execution may be interrupted by the scheduler and re-run “on-demand” VMs.
3. **high**: jobs are executed on “on-demand” VM and cannot be interrupted by the scheduler.

For our experiments we used the normal priority. In practice, we found that jobs that run on large machines (such as “mem3_ssd1_v2_x32”) are interrupted and executed “on-demand” after ~35 minutes of computation. For jobs who run on both “spot” and “on-demand” VM, the system charges both the time spent on the “spot” VM (and the interrupted) and the time spent on the “on-demand” VM (a new full execution). We report running times and costs only of the instance that successfully succeeded. The costs are obtained by multiplying running time reported by the Unix /usr/bin/time command, multiplied by the price of the VM used according to the instance used (**Supplementary Table 5-7**).

### **Supplementary Note 2.** Performance of low-coverage imputation methods

#### S2.1 Introduction

Generally, performing genotype imputation of lcWGS data is a computationally expensive task, as the running time of imputation methods depends on the number of samples and variants in the reference panel. We recently introduced GLIMPSE1[12], a low-coverage imputation method designed for large reference panels, that proposes a full haploid imputation model. This, together with other methodological advances, such as the use of the Positional Burrows Wheeler Transform[15], led to a reduction of running time for large reference panels. However, as whole-genome sequenced reference panels increase in size, the number of rare variants grows considerably, potentially leading to prohibitive running times. To overcome this major limitation, we introduce GLIMPSE2, which leverages the sparsity of reference panels at rare variants to perform efficient genotype imputation. In GLIMPSE2, each target sample is sparsely imputed and phased using the Li and Stephens model from a custom reference panel chosen using sparse PBWT matching (**Supplementary Note 1**).

#### S2.2 Methods

To assess the performance of GLIMPSE2, we first extracted 10,000 British samples from the UK Biobank whole-genome sequencing dataset, chosen to be fully unrelated (in-between them and to the reference panel). We used the remaining 140,199 samples as the reference panel with 13,296,637 markers on chromosome 20. Due to the heavy computational requirements of other methods, for this benchmark we only used 100 British samples as target for imputation and downsampled the reads at 0.1x, 1.0x and 4.0x coverage (**Supplementary Note 1**, results on the full 10,000 samples with GLIMPSE2 are shown in **Supplementary Note 6-7**).

We compared the performance of GLIMPSE2 to recently released lcWGS imputation methods: GLIMPSE v.1.1.1 (GLIMPSE1)[12] and QUILT v1.0.4[20]. We assessed imputation accuracy by measuring the squared Pearson correlation between imputed genotypes and high-coverage data within minor allele frequency (MAF) bins defined by the reference panel for the appropriate populations. As only

GLIMPSE2 fully supports small indels, we only focused on SNPs for the comparison between methods (for imputation accuracy of GLIMPSE2 at indels, see **Supplementary Note 7**).

We also evaluated the scaling properties of imputation methods using simulated data for European reference panels of 2,500, 10,000, 100,000, and 1,000,000 samples generated using MSPRIME v1.0[21] in a 10Mb region (**Supplementary Table 1**), similarly to what was shown with the IMPUTE5 method[13]. We simulated 1.0x coverage reads for 1,000 target samples from their haplotypes, by fixing the read length to 100 bp, a base quality of 30 and a sequencing error rate of 0.001. We then performed imputation using 1 Mb regions for GLIMPSE1 and QUILT v1.0.4 and 3.33 Mb for GLIMPSE2 and looked at their computational cost. QUILT v1.0.4 could not run on the reference panel containing one million samples due to memory constraints (>250 Gb for both QUILT_preparere_reference.R and QUILT.R scripts).

### S2.3 Results

As expected, imputation accuracy using the UK Biobank reference panel for British samples outperforms all other panels, both at rare and common variants (**Figure 1a**, **Supplementary Figure 2-4, Supplementary Table 2-4**). Regarding methods, GLIMPSE2 performs overall better than GLIMPSE1, mainly at rare variants, with the biggest difference at 0.1x coverage data (**Supplementary Figure 4**, **Supplementary Table 2-4**), but also at common variants (GLIMPSE2 imputation $r^2$=0.93; GLIMPSE1 imputation $r^2$=0.82). The improvements of GLIMPSE2 are also visible when compared to QUILT v1.0.4, which leverages the full reference panel and raw sequencing reads[20]. Indeed, we find that the two methods show similar results across all tested reference panels and coverages (**Figure 1a**, **Supplementary Figure 2-4, Supplementary Table 2-4)**.

Importantly, we also find that the GLIMPSE2 imputation costs are orders of magnitude smaller than for other methods (**Figure 1b, Supplementary Figure 5, Supplementary Table 5-7**). Assuming that all computing is performed “on-demand”, GLIMPSE2, GLIMPSE1 and QUILT v1.0.4 require 0.69, 12.21 and 485.60 British pounds of computing costs (as of November 2022), respectively, to impute 100 samples of 1.0x coverage on chromosome 20, which approximately corresponds to 0.35, 6.11 and 242.80 British pounds, for the entire genome of a single target sample (**Supplementary Table 7**). However, these are conservative estimates, as GLIMPSE1 and GLIMPSE2 allow imputation with shorter running times and smaller VMs, therefore avoiding the need for “on-demand” instances and achieve imputation of a whole genome for 1.1 and 0.08 British pounds per sample, respectively (**Supplementary Table 7**).

GLIMPSE is a method designed for many target samples. Indeed, the computational cost per sample of GLIMPSE2 is at maximum when imputing a single sample and decreases quickly by increasing the number of samples to impute until fixed costs are amortised (mainly sparse PBWT computations), and then it remains approximately linear with the increasing number of target samples (> 50 target samples, **Supplementary Figure 6c**). However, QUILT v1.0.4 is specifically designed to impute a single sample at a time. When we impute a single sample from the UKB reference panel, we find that the running time per sample on chromosome 20 was 1.4h, 40h and 16h for GLIMPSE2, GLIMPSE1 and QUILT v1.0.4, respectively, with an estimated whole-genome cost per sample of 0.92£, 26.40£ and 42.24£ on the UKB RAP (GLIMPSE1 shows increased running time, but less cost that QUILT due to QUILT's memory usage).

We investigate in detail the scaling properties of GLIMPSE2 by downsampling the reference panel on chromosome 20 into subsets of growing size (keeping only variant sites) and show that the running time of GLIMPSE2 increases sublinearly with reference panel size for every sequencing coverage, due to the PBWT selection and sparse imputation at rare variants. Indeed, the running time to impute a single variant decreases when increasing the reference panel size, due to the fact that most of the new variants brought from bigger reference panels are rare (**Supplementary Figure 6a**). This leads to a two-fold increase in running time per sample moving from a reference panel containing 2,500 samples and ~1.5 million variants to a reference panel containing 140,000 samples and ~12 million variants (**Supplementary Figure 6b**). Additionally, using simulated data, we empirically evaluated the model scaling of imputation methods up to a reference panel containing 1 million samples. We find that the running time to impute a single variant with QUILT v1.0.4 and GLIMPSE1 increases with reference panel size, while the running time of GLIMPSE2 decreases (**Supplementary Figure 7a**). Importantly, only GLIMPSE1 and GLIMPSE2 can run on the reference panel containing 2 million haplotypes, although with different memory usage (for GLIMPSE1, 87Gb per 1Mb chunk, for GLIMPSE2, 5.5Gb per 3.33 Mb chunk). Even in this case, we confirmed GLIMPSE2 has a running time per imputed variant that decreases with reference panel size, leading to a running time that is approximately increased by a factor of two when moving from a reference panel containing 2,500 samples and 100,000 variants to a reference panel containing 1 million samples and 2.2 million variants (**Supplementary Figure 7b**). These results allow us to conclude that the scaling properties of GLIMPSE2 are confirmed in both real data and simulation experiments.

### **Supplementary Note 3.** Imputation performance of low-coverage and SNP arrays

#### S3.1 Introduction

In the field of statistical genetics, many studies have been carried out using SNP arrays and imputed against a large reference panel[1]. We already showed the benefits of lcWGS compared to SNP array in a previous study for a wide range of applications[12], and other studies showed the improved accuracy of lcWGS compared to SNP-array imputation[20,22,23]. However, most of these studies did not have access to real SNP-array data and relied on simulated SNP arrays from high-coverage WGS.

Here, we use the UKB Axiom array data, a SNP array specifically designed to genotype the British population and to improve genotype imputation[1], and the recent WGS data from 150,119 samples[2] to benchmark low-coverage and SNP-array imputation in a more realistic scenario.

#### S3.2 Methods

We used UKB Axiom array data for 10,000 British samples used on chromosome 1, applied the same quality control performed in the original data release (UK Biobank Resource 531) and lifted over the data to the GRCh38 human reference genome, discarding strand flips (99.8% of the original variants maintained after the liftover). We then imputed SNP-array data using the UKB reference panel with BEAGLE v5.4 (release July 2022)[24] allowing pre-phasing from the reference panel. Additionally, we downsampled to five different coverages (0.1x, 0.25x, 0.5x, 1.0x and 4.0x) and imputed the data using GLIMPSE2.

#### S3.3 Results

Imputation of the UK Biobank Axiom array is quantitatively similar to lcWGS imputation of 0.25x data and 0.5x data, in agreement to our previous findings with simulated SNP arrays[12] (**Figure 1c**). Higher sequencing coverages, such as 1.0x and 4.0x data, confidently outperform the Axiom array with the biggest difference seen at rare variants (for 1.0x and 4.0x coverage, accuracy improvement of $r^2 > 0.1$ and $r^2 > 0.17$ for variants with a MAF < 0.01%, **Figure 1c**). It is important to note here that this represents an ideal scenario for SNP array imputation, as the UK Biobank Axiom array has been specifically designed for this dataset to improve imputation, it matches the ancestry of the reference panel, and it does not suffer from ascertainment bias for this population.

SNP-array imputation is computationally more efficient than lcWGS imputation, with a whole genome imputation cost that is less than one cent per sample. This is at least one order of magnitude cheaper than lcWGS imputation with GLIMPSE2. However, GLIMPSE2 remains economically sustainable and requires similar hardware requirements as SNP-array imputation methods.

We investigate more in detail the consequences of these results for downstream applications in **Supplementary Note 6-7**, where we look at the impact in GWAS and rare variant burden testing.

### **Supplementary Note 4.** Imputation of Simons Genome Diversity Project dataset

#### S4.1 Introduction

We show in **Supplementary Note S2** that the UKB represents a valuable resource as a reference panel for imputation for the British population, providing better accuracy than other reference panels such as the HRC[9] and KGP[6]. However, the UKB reference panel does not offer a global representation of human diversity, and for non-European populations it is unclear whether the UKB is beneficial compared to the KGP reference panel. Indeed, the KGP offers an extremely valuable and diverse resource covering many world-wide ethnicities and ancestries (**Supplementary Note S1.1.2**). Here, we aim to evaluate the imputation performance of the UKB and KGP reference panels, across a wide range of populations. To do so, we use publicly available data from the Simons Genome Diversity Project (SGDP)[25] (**Supplementary Note 1.1.4**) and compare imputation performance from the two reference panels.

#### S4.2 Methods

We imputed 276 SGDP individuals from 129 world-wide populations downsampled at 1.0x coverage for chromosome 20, using the UKB and KGP reference panels. Prior to imputation, we removed samples from KGP in common with SGDP. We used high-coverage data for SGDP (**Supplementary Note 1.4**) mapped and genotyped using a customised procedure that was optimised for population genetic analysis[10] and assessed imputation accuracy at the intersection of the biallelic SNPs present in UKB and KGP. As we do not have a precise estimate of minor allele frequencies, we look at the non-reference discordance rate (NRD) between the two reference panels for each sample.

#### S4.3 Results

Overall, imputation performances follow the target population representation in the reference panel, as expected. The UK Biobank reference panel substantially improves the imputation of European individuals, especially those with Northern European ancestry (**Supplementary Figure 8a)**. In particular, the biggest difference between panels is found for the Orcadian, and English samples (NRD reduction > 74%), followed by Norwegian and Icelandic samples (NRD reduction > 55%) (**Supplementary Figure 8b)**. Significant improvements for non-European populations are found for the Maori genome from New Zealand (NRD reduction=62%), a sample with large components of European and Polynesian ancestries[25] , the latter is not well represented in KGP. The UKB reference panel performs overall better for South Indian samples, mainly for samples from Pakistan (Balochi, Brahui, Burusho, Hazara, Kalash, Makrani, Pathan, and Sindhi; NRD reduction > 14%), although smaller improvements are seen for Punjabi, due to their presence in KGP. On the other hand, KGP outperforms UKB reference panels for individuals of Native American and East Asian ancestries. For African ancestry, we find overall better imputation with UKB except for populations represented in KGP (Mandinka, Mende, and Yoruba), but we still see higher NRD values compared to other populations in absolute terms (**Supplementary Figure 9**), likely due to underrepresentation and larger genetic diversity of African populations compared to other populations across the world[26].

### **Supplementary Note 5.** Imputation of ancient European populations

#### S5.1 Introduction

Postmortem damage leads to high levels of missingness in the vast majority of ancient genomes. In a study we recently released, we showed that imputation can improve low-coverage ancient DNA data for most ancestries[27]. In that study, we used KGP as a reference panel. Given the imputation performance of present-day European individuals using the UK Biobank reference panel, here we investigate if this increased accuracy can also be seen in ancient DNA samples. As before, we downsampled high-coverage ancient genomes to low coverage and imputed these with GLIMPSE2 and varying reference panels, i.e., KGP vs. UKB. We then used the high-coverage data to validate the imputation results. For that, we selected four publicly available ancient genomes with coverage of at least 18x from relevant populations according to **Supplementary FIgure 8-9**: an ancient Viking individual from Iceland (HSJ-A-1[28], ~1,000 years before present (ybp), 29x coverage), an ancient Neolithic individual sample from Hungary (NE1[29], European farmer from Hungary, ~7150 ybp, 18x coverage), an European Mesolithic individual (Loschbour[30], Western Hunter-Gatherer from Luxembourg, 18x coverage), and an Early Bronze Age individual sample from Kazakhstan (Yamnaya[31], Yamnaya from Kazakhstan, 26x coverage). The ancestries of these last three individuals are present in most present-day Europeans with variable proportions across the continent[30,32]. Southern Europeans carry more Early European Farmer ancestry than their Northern counterparts, and the opposite pattern is seen with Western Hunter-Gatherer and Yamnaya ancestries.

#### S5.2 Methods

The publicly available BAM files were aligned to the hg19 reference genome. To use the UKB as a reference panel for imputation, we re-mapped the data to the GRCh38 reference genome. For that, we extracted the fastq files from the original BAM files using the command bedtools bamtofastq from bedtools v2.29.2[33]. We then aligned the data to the GRCh38 reference genome using mapache[34], a Snakemake utility pipeline designed to map ancient DNA sequences, with default parameters.

To estimate imputation accuracy, we generated a validation dataset from the high-coverage BAM files. Due to postmortem damage, ancient DNA sequences contain more errors than their present-day counterparts. Therefore, to minimise the impact of such errors on the genotype calls, we followed the same approach as in Moreno-Mayar et al.[35]: i) genotype calling with BCFtools[36] v1.12 using reads with a minimum of 30 mapping quality (-q 30), keeping only bases with quality of at least 20 (-Q 20), and with the parameter -C 50, as suggested by the SAMtools[36] developers when handling data mapped with BWA that contain more mismatches; ii) keeping exclusively the sites present in the 1000 Genomes accessible genome strict mask[37]; iii) exclusion of sites inside repeat regions (RepeatMask regions in UCSC Table Browser[38]; iv) removal of sites that are outliers regarding depth: either below the maximum of one third of the mean depth of coverage (DoC) and 8x (max(1/3DoC, 8), and depth above twice the DoC; v) removal of sites with QUAL field below 30.

We restricted the analyses to chromosome 20. We used SAMtools v1.12 to downsample the BAM files to several depths of coverage (0.1x, 0.25x, 0.5x, 1.0x, 4.0x). We proceeded to impute the data as previously described.

Finally, we assessed imputation accuracy at the intersection of the biallelic SNPs (no singletons) present in UKB and KGP (**Supplementary Note S1.1.1-S1.1.2**). We generated the intersection with BCFtools.

#### S5.3 Results

Imputation results for all the reference panels for the downsampled data are shown in (**Supplementary FIgure 10**). Compared to KGP, the UKB reference panel substantially improves imputation for HSJ-A-1, Loschbour and Yamnaya samples, both at common and rare heterozygous sites. For the Viking genome (HSJ-A-1), these results agree with our previous finding that the UKB reference panel leads to a boost in imputation accuracy in Northern European individuals. We find that the UK Biobank reference panel is particularly suitable to impute the Loschbour and Yamnaya genomes, whose ancestries are present to a greater extent in Northern Europeans, including the British[30,32]. In the case of NE1, we also find that the UK Biobank increases imputation accuracy, but to a lesser degree at very rare variants (0.1%<MAF<1%) and coverages below 0.5x.

The results show that it is possible to recover rare variants (MAF < 0.1%) from ancient DNA samples of these ancestries (non-reference concordance > 60%), despite pervasive post-mortem damage, such as fragmentation and deamination[39].

### **Supplementary Note 6.** Common-variant association analyses

#### S6.1 Introduction

Low-coverage WGS has been shown to be a reliable technology for functional analysis of complex traits and particularly for expression quantitative trait loci[40], burden test[14], genome-wide association studies (GWAS)[41–43] and polygenic risk score analysis[22]. Here we evaluate the performance of genome-wide association scans using imputed low-coverage WGS and SNP-array data across a wide range of phenotypes, comparing these with association scans performed with high-coverage WGS data. This approach is particularly relevant on the UK Biobank data as it contains both WGS and SNP-array data and hundreds of phenotypes, and therefore allows assessing the respective performance of lcWGS and SNP array to perform GWAS at realistic scale.

#### S6.2 Methods

We used chromosome 1 for a subset of 10,000 UK Biobank individuals of white British ancestry randomly sampled and a total of 99 phenotypes, selected as phenotypes with less than 10% of missing data in our call set across anthropomorphic traits and blood measurements. We used plink2[44] with default parameters and sex, age and the first 10 PCs as covariates to test phenotypes for associations with seven different call sets: the high-coverage WGS, five low-coverage WGS (0.1x, 0.25x, 0.5x, 1.0x, 4.0x) and the UKB Axiom array. We selected (i) genome-wide significant loci (p-value < $5e^{-08}$) and (ii) independent, being at least 500 kb apart.

To assess the accuracy of GWAS performed using imputed call sets, we first compared association strength (i.e p-values) and effect sizes (i.e betas) to the high coverage GWAS. For this, we computed the Pearson correlation ($r^2$) between p-values and betas of each GWAS performed using imputed callsets and the high coverage GWAS.

We then assess the ability of GWAS performed using imputed low-coverages and SNP array to distinguish significant from non-significant signals, considering the high-coverage GWAS to be the ground truth. For this, we computed the sensitivity (i.e true positive rate, which is the proportion of genome-wide significant associations that can be retrieved) and specificity (i.e true negative rate, which is the proportion of genome-wide non-significant associations that can be retrieved) of GWAS performed using imputed callsets.

#### S6.3 Results

We find a total of 22 phenotypes with at least one genome-wide significant locus in the validation GWAS. When testing these phenotypes using imputed call sets (**Supplementary Figure 11-13**, for 0.1x, 0.25x, 1.0x and Axiom array across 6 selected phenotypes), we find that increasing the coverage increases the accuracy of association scans, as expected, with $r^2$ ranging from 0.92 to 0.99 for betas (for 0.1x and 4.0x, respectively), and from 0.91 to 0.99 for p-values (**Figure 1d**). This highlights that even with 0.1x coverage, the majority of GWAS signals have similar effect sizes and association strength with the phenotype. We observe that 0.25x sequencing has a similar accuracy than the UK Biobank Axiom array (betas $r^2$=0.97 and 0.96; p-values $r^2$=0.97 and 0.97, for 0.25x and Axiom array, respectively). A coverage of 1x leads to better p-value and effect size estimates than the Axiom array, although this SNP array has been specifically designed for the British population.

Moreover, we find that the accuracy of association strength and effect sizes is similar. However, imputed 0.25x-0.5x call sets allow retrieving more true positive signals compared to the Axiom array (**Supplementary Figure 14a**). In addition, we observe that specificity was higher than 99.99% for all imputed call sets (**Supplementary Figure 14b**), which shows that GWAS based on imputed call sets does not lead to an excess of false negatives. Together, these results highlight that reducing the sequencing depth has a relatively small impact on the discovery power and accuracy of downstream GWAS.

### **Supplementary Note 7.** Rare-variant association analyses

#### S7.1 Introduction

Functional analysis of rare variants plays a significant role in Mendelian conditions as well as in many complex traits. Pivotal studies are currently discovering and refining the functional properties of variants in a large whole-exome and whole-genome sequencing cohorts, identifying predicted loss-of-function variants and improving the knowledge of the impact of rare variants in many diseases[45,46]. Additionally, recent advances in haplotype phasing allow the examination of loss-of-function compound heterozygous variants on complex traits[5], complementing current knowledge of gene essentiality in the human genome. While high-coverage WGS remains crucial to make new discoveries, lcWGS imputation from large reference panels can offer a valid alternative, due to the improved accuracy at rare variants compared to SNP arrays. Here, we measure imputation performance at INDELs compared to high coverage data and we investigate the loss of power for burden-test functional analysis from lcWGS and SNP array data at rare functional coding variants.

#### S7.2 Methods

In order to assess imputation accuracy at rare variants of lcWGS and SNP-array data, we used 10,000 British samples with both 1.0x-coverage and SNP-array data, which we imputed genome-wide using GLIMPSE2 and BEAGLE v5.4 (release July 2022)[24], respectively. First, we assessed the genome-wide imputation performance at INDEL sites, by computing a genome-wide imputation $r^2$ aggregating across the 10,000 samples on chromosome 1. For this, we used the data presented in **Supplementary Note 3** and **6**. Second, we assessed the respective imputation accuracy of loss-of-function (LoF), missense and synonymous variants genome-wide at the gene level, partitioned using Genebass annotations (based on Ensembl VEP and LOFTEE[45]). For this, we measured concordance as the squared Pearson correlation ($r^2$) between the imputed and high-coverage allele dosages of rare variants (MAC < 200 in high-coverage data) for each class of variant annotation separately, across all protein-coding genes. Measuring $r^2$ has the advantage of quantifying the loss of power in statistical testing. We did not consider genes for which we could not compute an $r^2$ coefficient, looking at more than 80% of the original LoF genes, and more than 96% of the original missense and synonymous genes.

#### S7.3 Results

As expected, imputation accuracy at INDELs (**Supplementary Figure 15a**) is lower than what we report for SNPs (**Figure 1c**), for both low-coverage and SNP array imputed data. However, the results show a similar trend. Coverages such as 0.25x and 0.5x perform similarly to the Axiom array, and coverages starting from 1.0x perform better, mainly at rare variants (MAF<0.1%, **Supplementary Figure 15a**). Notably, other low-coverage imputation methods seemingly cannot perform imputation of INDELs, as it can potentially harm imputation accuracy at the neighbouring SNPs. However, GLIMPSE2 is able to impute INDELs with a model designed for complex genetic variants (using a haplotype scaffold, **Supplementary Note S1.2.2.6**).

We find that imputed data at 1.0x coverage significantly outperforms the Axiom array also in burden test analysis for each annotation used (Wilcoxon non-parametric test P-value < 2e-16, **Supplementary Figure 15b**), confirming what we showed in our previous study[12]. Furthermore, we note that missense and synonymous variants are better imputed than LoF, likely resulting from the low allele frequency of LoF due to their strong negative selection and presence of INDELs. Overall, these results suggest that both low-coverage and SNP array imputation data from a large reference panel constitute a promising approach for burden-test analysis with a moderate loss of power. However, lcWGS at a coverage of 1.0x outperformed population-specific SNP arrays for this analysis.

### Supplementary Figures

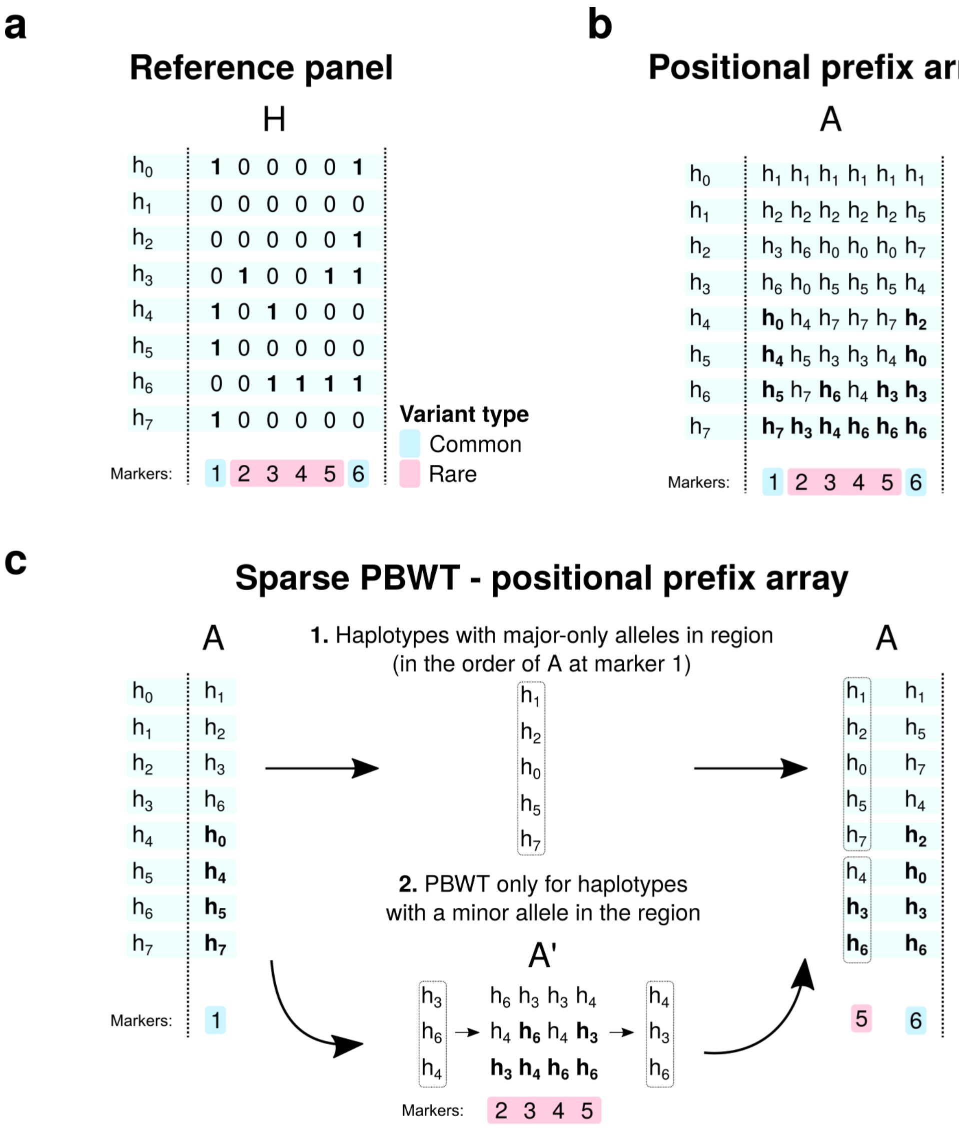


**Supplementary Figure 1: Sparse PBWT positional prefix array computation.** (**a**) We consider a reference panel $H$ with $M = 6$ markers and $N = 8$ haplotypes, $h_0, h_1, \dots, h_7$. Here, marker 1 and marker 6 are common variants (light blue), and markers from 2 to 5 are rare variants (red). (**b**) Full prefix array $A$ of the reference panel. (**c**) Sparse PBWT positional prefix array. At common variants (markers 1 and 6) the standard PBWT update is performed (light blue sites). At rare variants (red sites), no computation is required for the $L = 5$ haplotypes containing only the major allele in the region ($h_0, h_1, h_2, h_5, h_7$) and they can be copied at the beginning of $A_5$ in the same relative order as they appear in $A_1$. For the haplotypes that contain the minor allele in the region ($h_3, h_4, h_6$), we compute the positional prefix array $A'$ at the rare variants in the interval. The last positional prefix array ($A'_5$) can be directly copied into $A_5$ from position $N - L$.

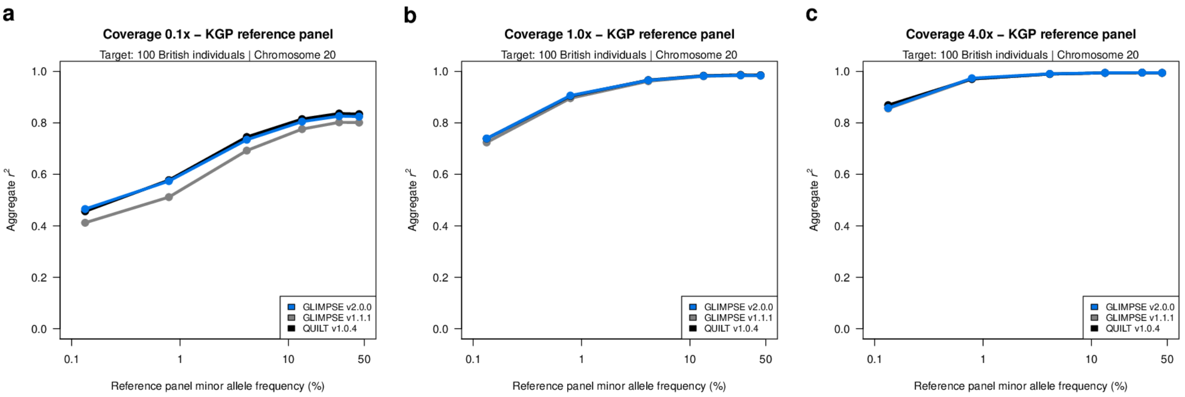


**Supplementary Figure 2: Imputation performance for the KGP reference panel.**
(**a**,**b**,**c**) Imputation accuracy (Aggregate $r^2$, y-axis) for 100 British samples at different sequencing coverage when using the KGP reference panel, imputed with GLIMPSE1 (grey) or QUILT v1.0.4 (black) and GLIMPSE2 (blue) for chromosome 20. Each panel represents a different sequencing coverage: (**a**) 0.1x, (**b**) 1.0x, and (**c**) 4.0x coverage. Accuracy in plotted against minor allele frequency of the reference panel (x-axis, log-scale).

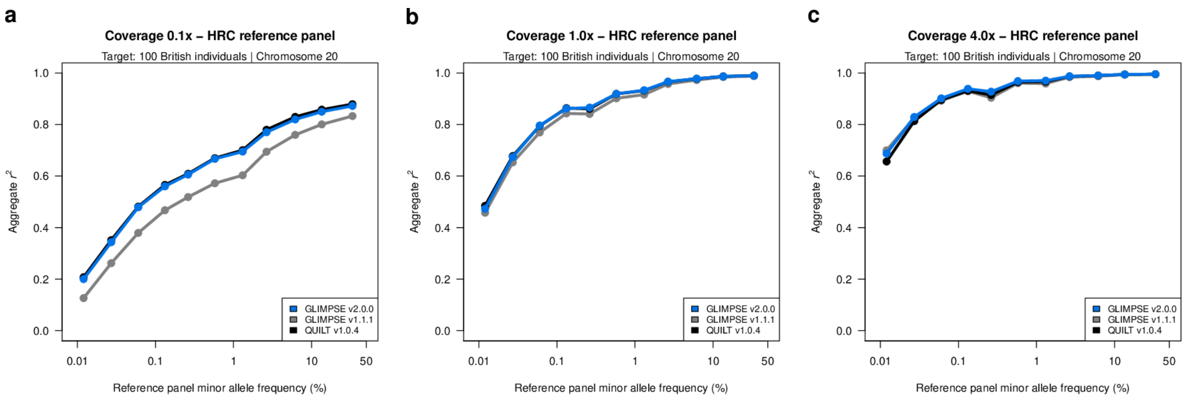


**Supplementary Figure 3: Imputation performance for the HRC reference panel.**
(**a,b,c**) Imputation accuracy (Aggregate $r^2$, y-axis) for 100 British samples at different sequencing coverage when using the HRC reference panel, imputed with GLIMPSE1 (grey) or QUILT v1.0.4 (black) and GLIMPSE2 (blue) for chromosome 20. Each panel represents a different sequencing coverage: (**a**) 0.1x, (**b**) 1.0x, and (**c**) 4.0x coverage. Accuracy in plotted against minor allele frequency of the reference panel (x-axis, log-scale).

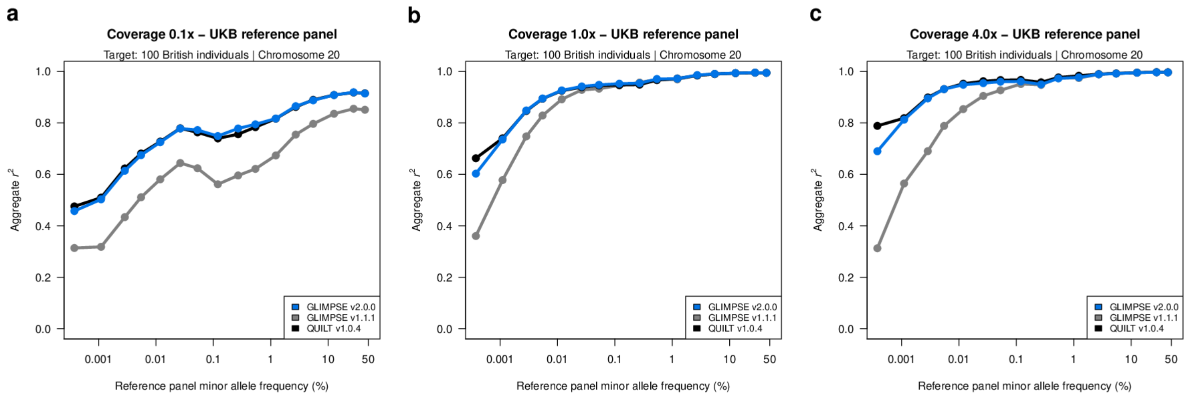


**Supplementary Figure 4: Imputation performance for the UKB reference panel.**
(**a**,**b**,**c**) Imputation accuracy (Aggregate $r^2$, y-axis) for 100 British samples at different sequencing coverage when using the UKB reference panel, imputed with GLIMPSE1 (grey) or QUILT v1.0.4 (black) and GLIMPSE2 (blue) for chromosome 20. Each panel represents a different sequencing coverage: (**a**) 0.1x, (**b**) 1.0x, and (**c**) 4.0x coverage. Accuracy in plotted against minor allele frequency of the reference panel (x-axis, log-scale).

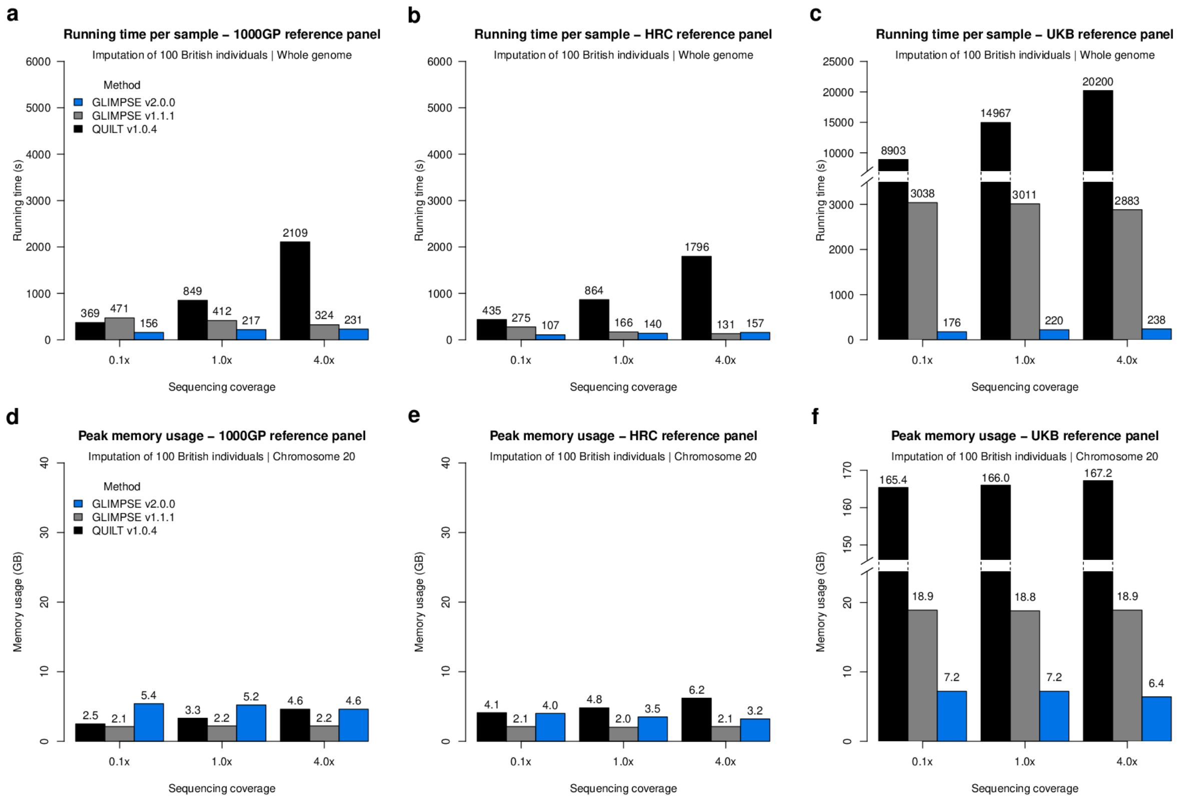


**Supplementary Figure 5: Computational performance of imputation methods using real reference panels.**

(**a-c**) Wall-clock time of imputation methods (in seconds, y-axis) at different sequencing coverages (0.1x, 1.0x and 4.0x, x-axis) for 100 British samples imputed with GLIMPSE1 (gray), QUILT v1.0.4 (black) and GLIMPSE2 (blue). Each panel represents a different reference panel. (**a**) KGP, (**b**) HRC, and (**c**) UKB. (**d-f**) Peak memory usage of imputation methods (in GB, y-axis) at different sequencing coverages (0.1x, 1.0x and 4.0x, x-axis) for 100 British samples imputed with GLIMPSE1 (gray) or QUILT v1.0.4 (black) and GLIMPSE2 (blue). Each panel represents a different reference panel: (**d**) KGP, (**e**) HRC, and (**f**) UKB.

All imputation jobs have been running using 4 threads/cores. GLIMPSE1 and QUILT were run in 2Mb regions. GLIMPSE2 was run in 4Mb regions. GLIMPSE2 and QUILT were run directly using BAM files, GLIMPSE1 was run on already computed genotype likelihoods (genotype likelihood computation is not included in the time).

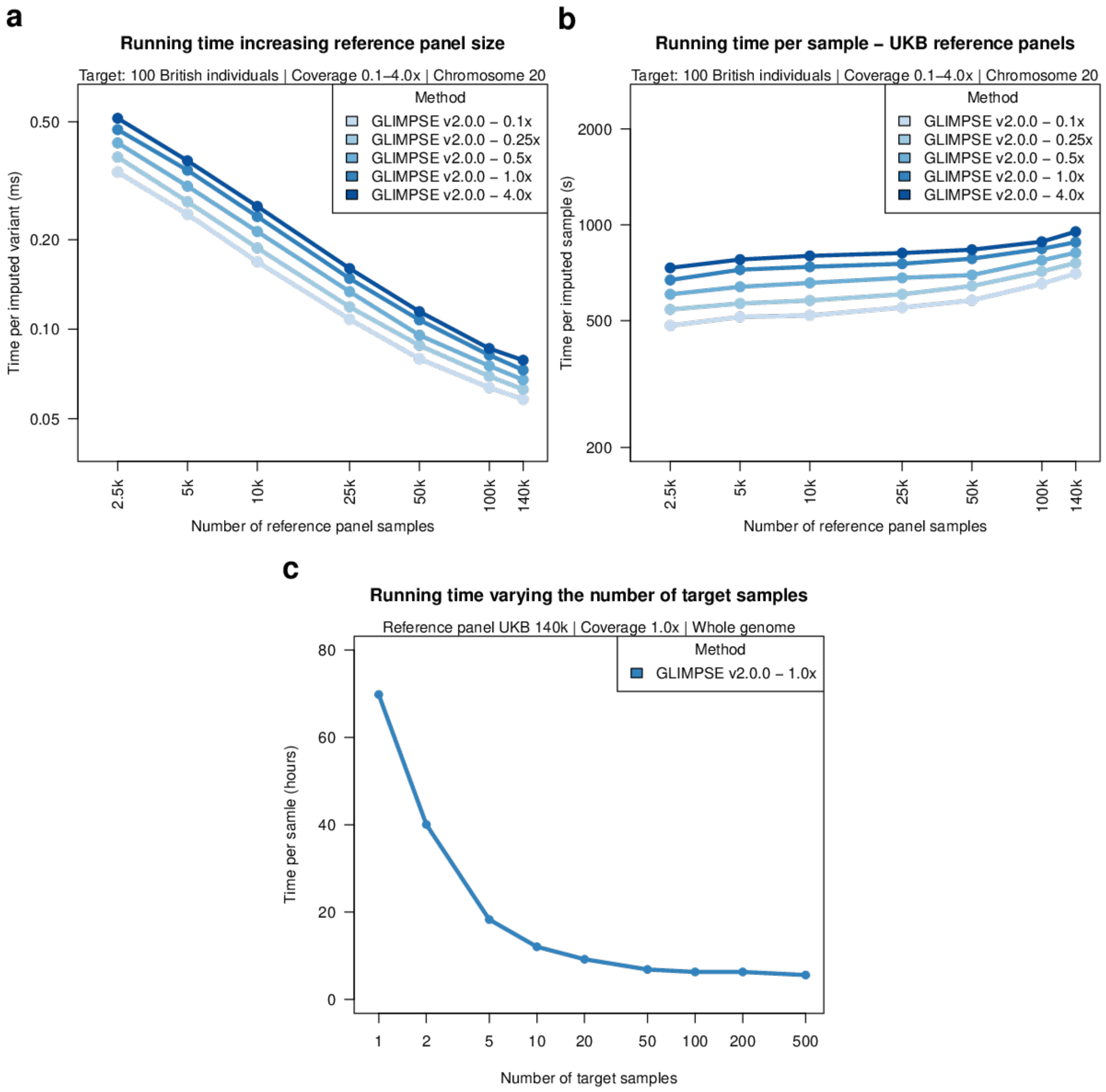


**Supplementary Figure 6. Model scaling of GLIMPSE2 using real data.**
(**a-b**) Log-log plots showing running time of GLIMPSE2 (run with 4 threads) to impute (**a**) a variant or (**b**) a sample (y-axis, log scale) on chromosome 20 using multiple coverages for 100 British samples (shades of blue) and multiple subsets of the reference panel (x-axis, log-scale), only retaining variable positions. (**c**) Running time (y-axis) varying the number of target samples (x-axis) using one thread with the full UKB reference panel for 1.0x coverage data.

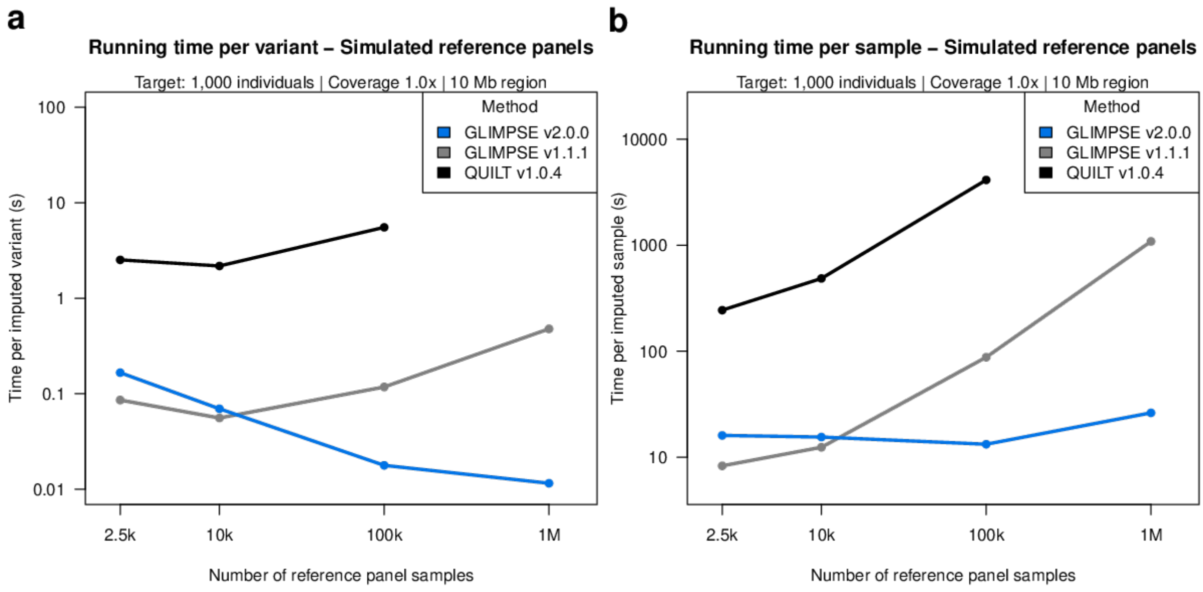


**Supplementary Figure 7. Model scaling using simulated data.**
(**a-b**) Log-log plots showing running time of GLIMPSE1 (gray), QUILT v1.0.4 (purple) and GLIMPSE2 v1.0.0 (blue) to impute (**a**) a variant or (**b**) a sample (y-axis, seconds, log scale) on a 10Mb region simulated with MSPRIME using 1.0x coverage simulated data and multiple subsets of the reference panel (x-axis, log-scale). The target dataset consisted of 1,000 samples and all methods were run on a single thread/core. GLIMPSE1 was run on already computed genotype likelihoods (genotype likelihood computation is not included in the time). QUILT v1.0.4 could not run on the reference panel containing one million samples due to memory constraints.

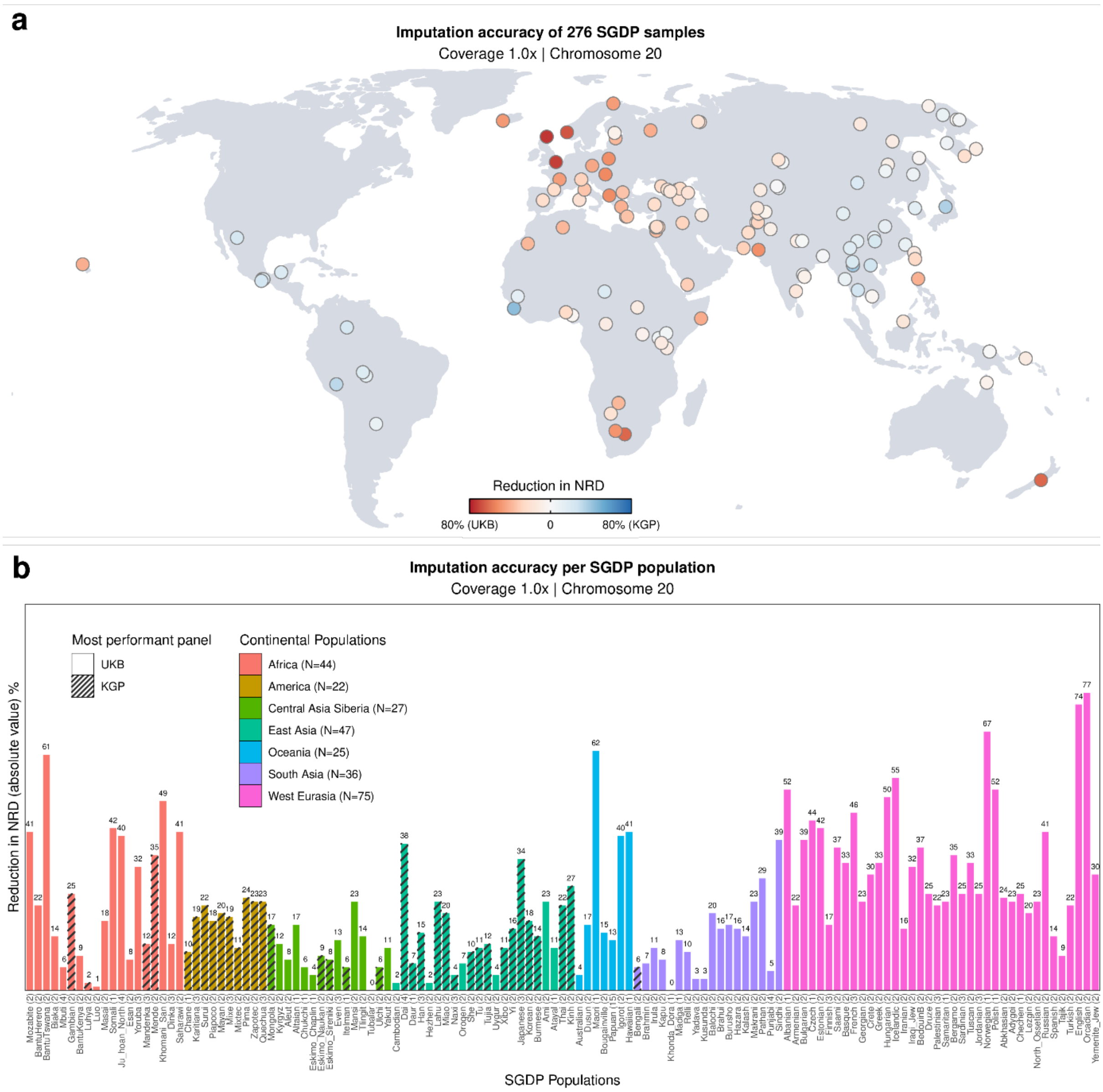


**Supplementary Figure 8: Performance of SGDP samples using different reference panels.** (**a-b**) Comparison between KGP and the UKB reference panels to impute 276 SGDP samples across 129 world-wide populations at 1.0x coverage on chromosome 20. (**a**) Per-sample comparison. Each circle represents one sample of SGDP and is coloured according to the reduction in NRD achieved when using the UKB reference panel (red) or KGP (blue). Location represents the geographical origin of the sample. (**b**) Population-level comparison. Samples belonging to the same population (x-axis) have been considered together (number shown in the x-axis label), showing the reduction of NRD between the two panels (y-axis). Populations have been coloured and ordered according to the continent and country of origin. Striped bars represent populations where KGP performs better than UKB reference panels.

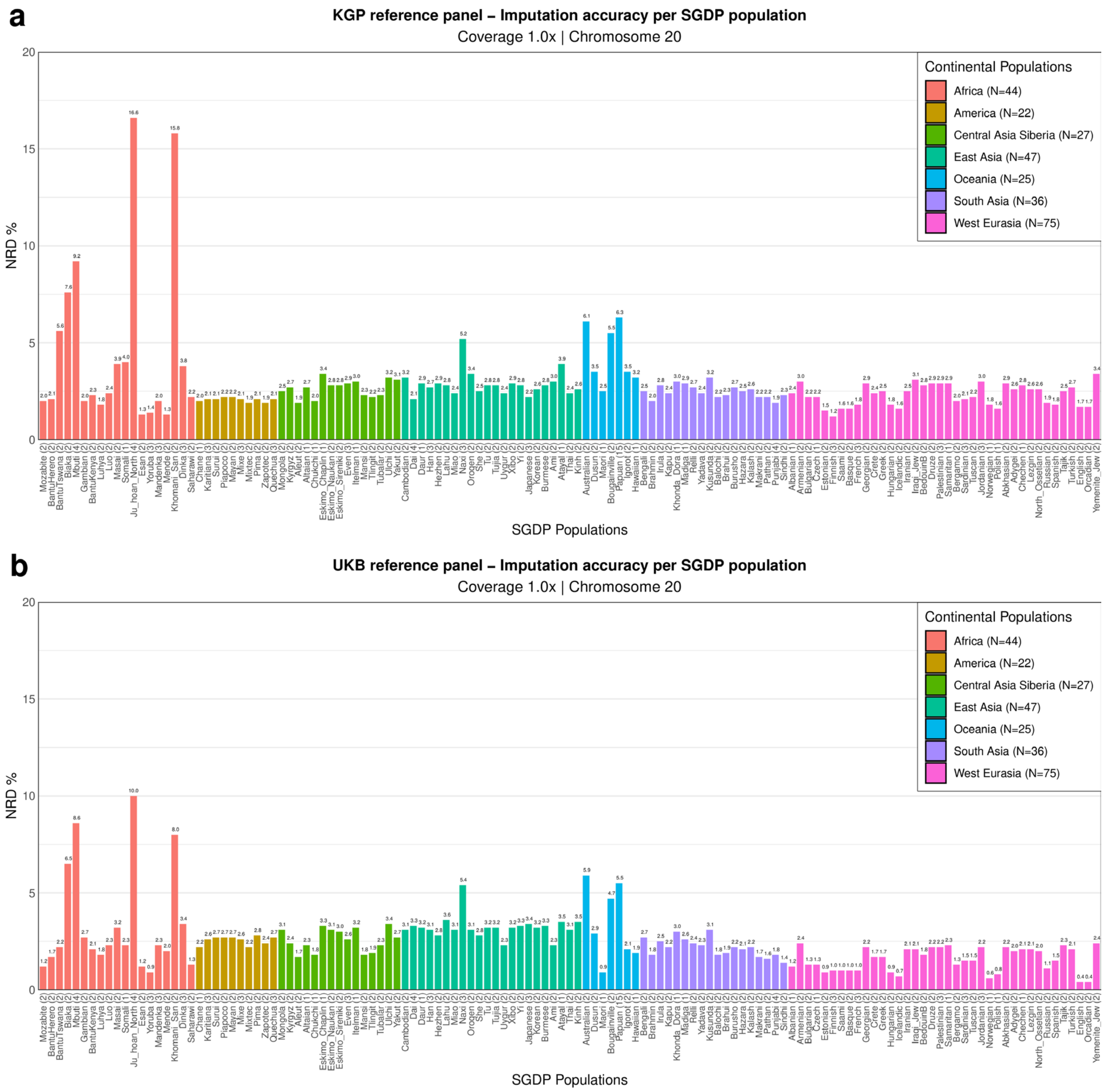


**Supplementary Figure 9: Performance of SGDP populations using different reference panels.** (**a-b**) Population-level comparison between KGP and the UKB reference panels to impute 276 SGDP samples across 129 world-wide populations at 1.0x coverage on chromosome 20. Samples belonging to the same population have been considered together, showing the NRD (y-axis). (**a**) KGP reference panel (**b**) UKB reference panel.

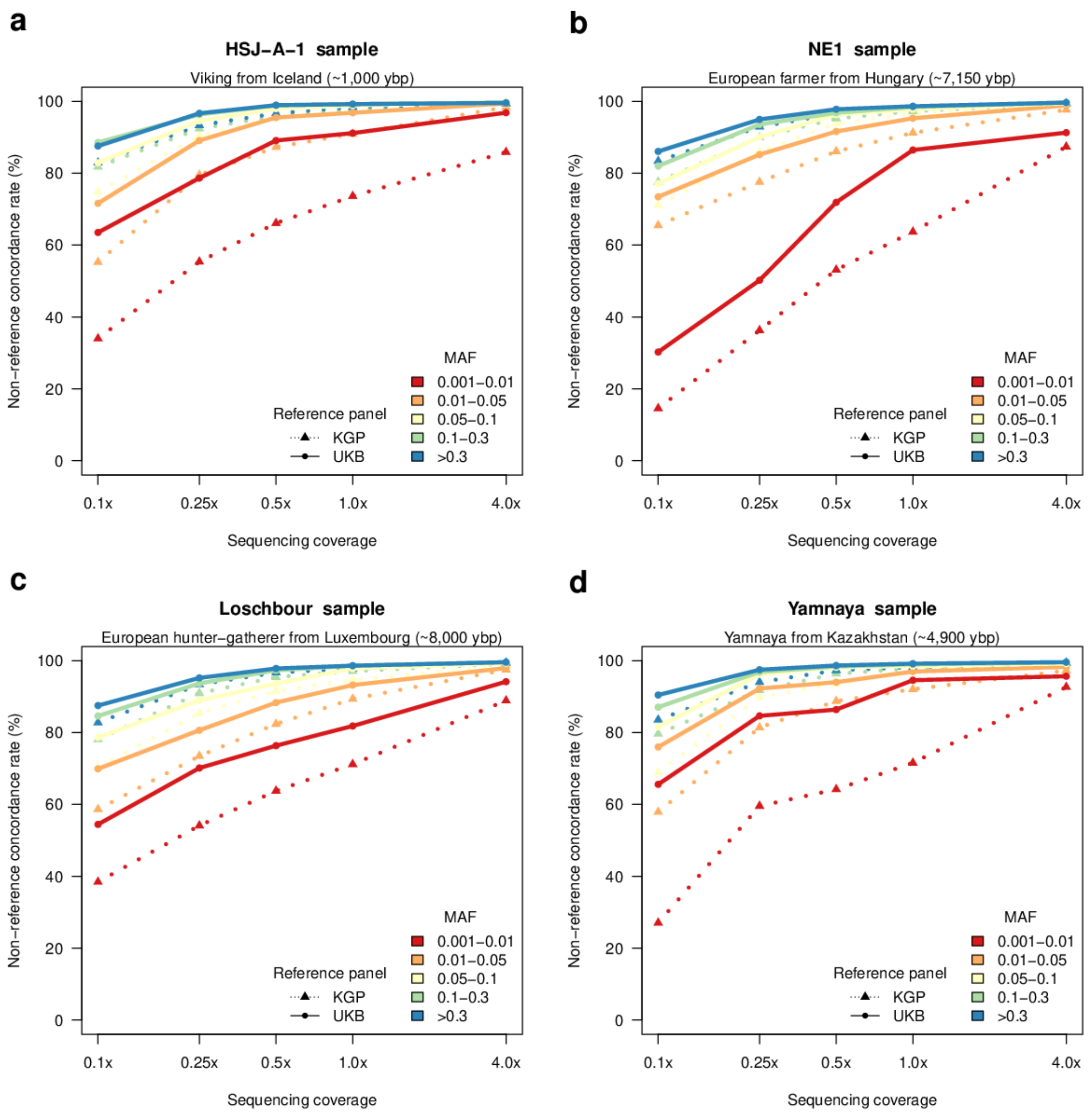


**Supplementary Figure 10: Imputation performance of ancient European samples using different reference panels.**

(**a-d**) Imputation accuracy (non-reference concordance rate percent, y-axis) of KGP (dashed line) and the UKB reference panel (solid line) to impute 4 ancient samples at different coverages (x-axis, log-scale) on chromosome 20. (**a**) HSJ-A-1, Viking sample, (**b**) NE1, European farmer, (**c**) Loschbour, European hunter-gatherer, (**d**) Yamnaya sample.

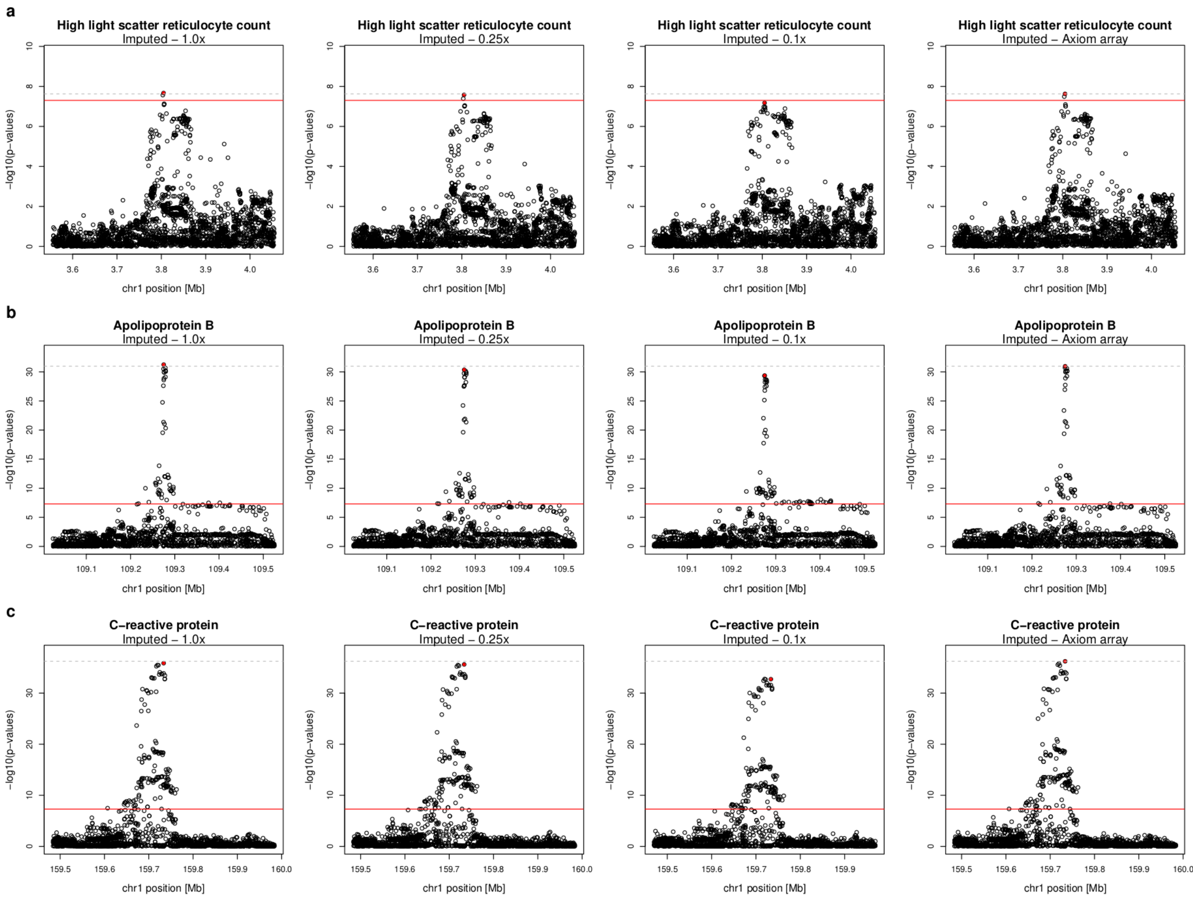


**Supplementary Figure 11. Genome-wide association of imputed call sets for high light-scatter reticulocyte count, apolipoprotein B and C−reactive protein phenotypes.**

GWAS scans for imputed 0.1x, 0.25x, 1.0x and Axiom array call sets across three selected phenotypes **(a-c)**. Association strength (-log10(p-values), y-axis) computed using plink2[44] along the chromosome 1 positions (x-axis). Red plain lines represent the Bonferroni genome-wide significance threshold at -log10(5e$^{-08}$); grey dotted lines show the association strength at the lead Axiom array hit; red dots show the lead variant in the validation GWAS.

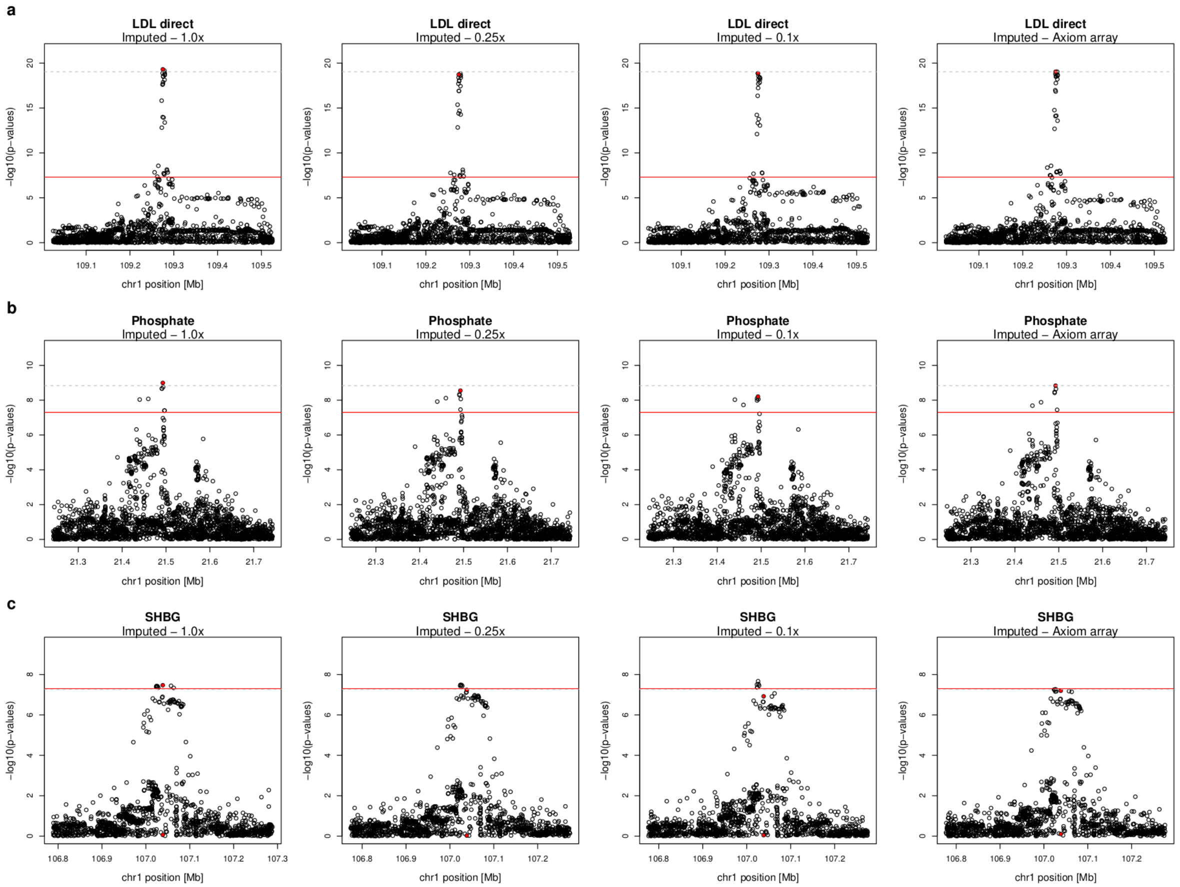


**Supplementary Figure 12. Genome-wide association of imputed call sets for direct LDL cholesterol, phosphate and sex hormone-binding globulin (SHBG) phenotypes.**

GWAS scans for imputed 0.1x, 0.25x, 1.0x and Axiom array call sets across three selected phenotypes **(a-c)**. Association strength (-log10(p-values), y-axis computed using plink2[44] along the chromosome 1 positions (x-axis). Red plain lines represent the Bonferroni genome-wide significance threshold at -log10($5e^{-08}$); grey dotted lines show the association strength at the lead Axiom array hit; red dots show the lead variant in the validation GWAS

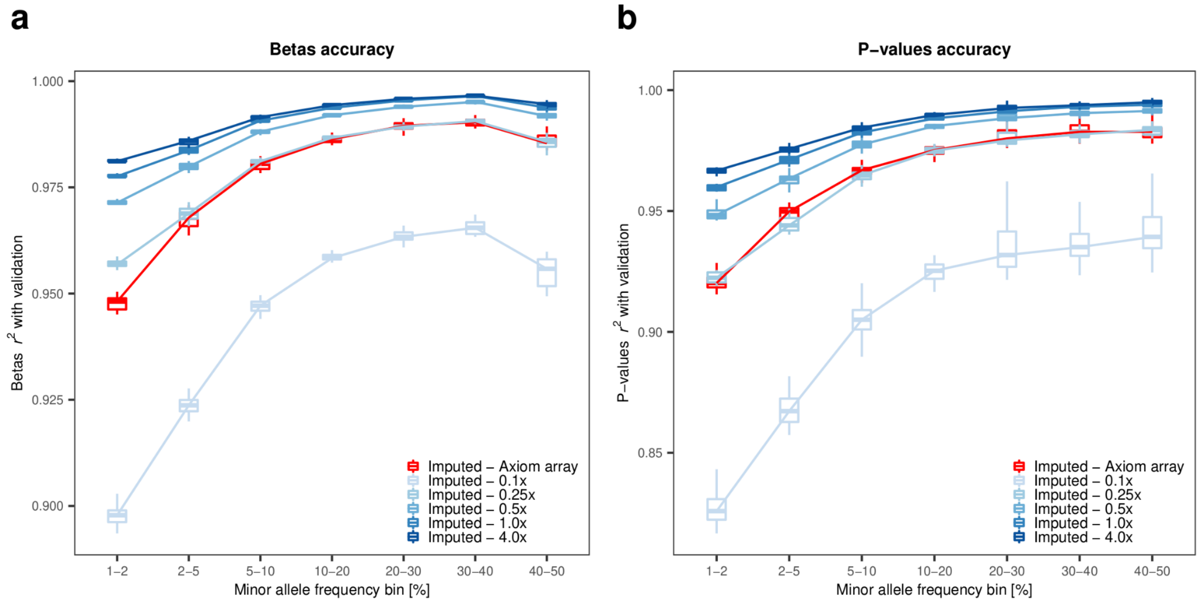


**Supplementary Figure 13. Accuracy of genome-wide association using imputed call sets.** (**a**) Squared Pearson correlation (y-axis) between the betas (effect sizes) computed from imputed and validation (high coverage) call sets, across the 22 phenotypes tested. (**b**) shows the same information for p-values. In both cases, the correlation is plotted as a function of minor allele frequency bins (x-axis).

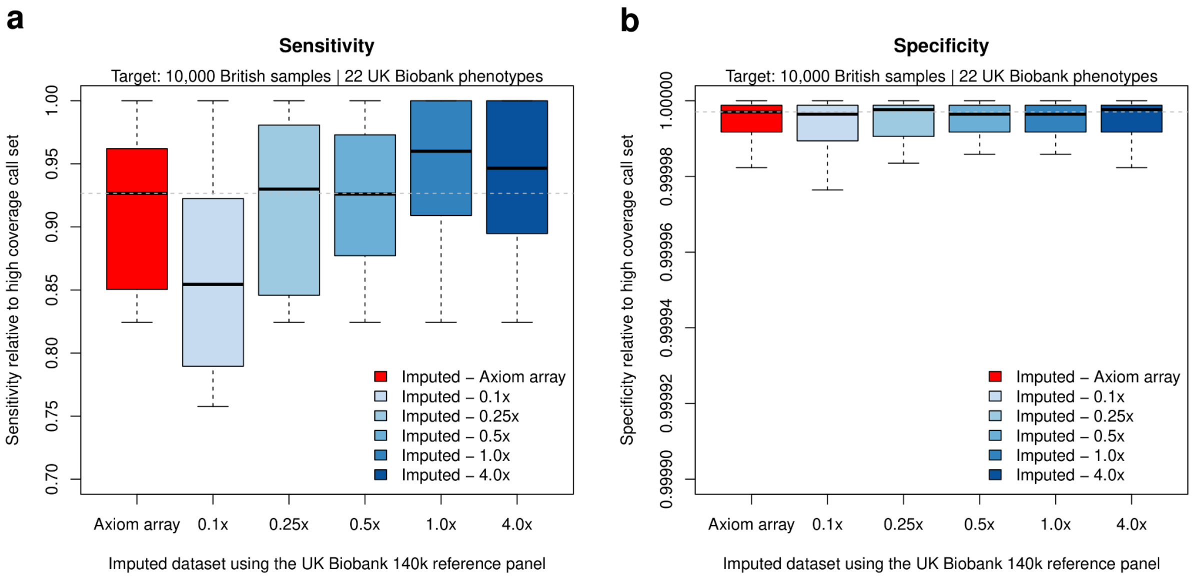


**Supplementary Figure 14. Sensitivity and specificity of genome-wide association using imputed call sets.** (**a-b**) Sensitivity (**a**, y-axis) and specificity (**b**, y-axis) of GWAS by comparing with the validation GWAS across the 22 phenotypes examined. The x-axis shows the different imputed call sets (0.1-4.0x, different shades of blue, GLIMPSE2 imputation; UKB Axiom array, red, imputed). Grey dotted lines represent the medians for GWAS using the Axiom array call set.

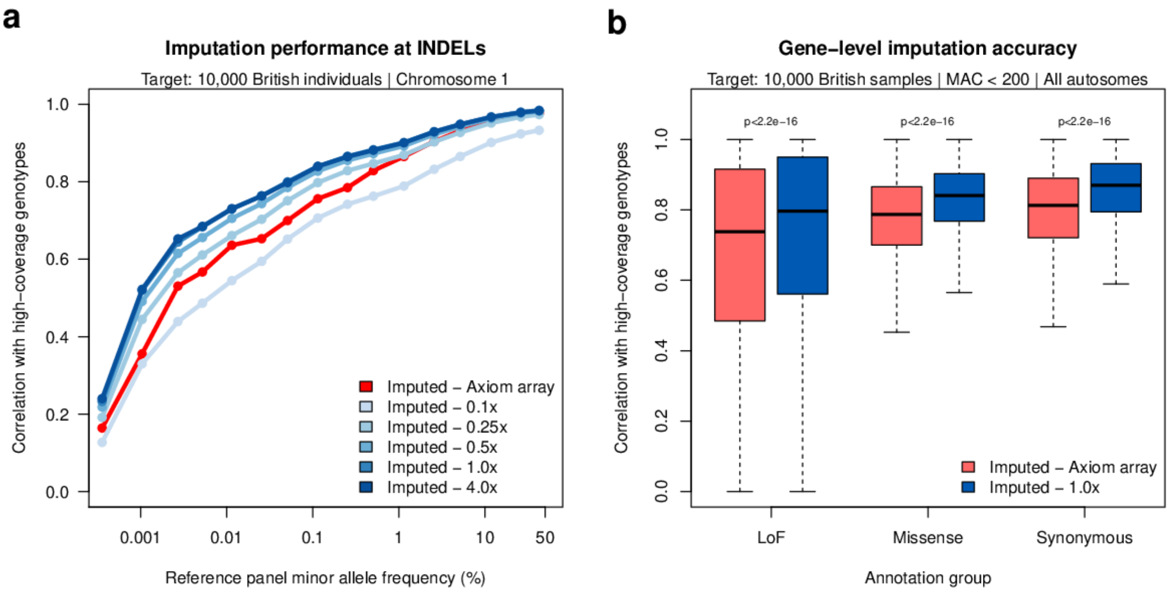


**Supplementary Figure 15. Performance at genomic annotations compared to high-coverage data.** (**a-b**) Imputation performance of 10,000 British samples imputed using the UKB reference panel across coverages (0.1-4.0x, different shades of blue, GLIMPSE2 imputation) and the UKB Axiom array data (red). (**a**) Imputation accuracy at INDEL sites. (**b**) Gene-level imputation accuracy ($r^2$, y-axis) at rare Genebass functionally annotated variants (LoF, loss of function; missense, synonymous variants; MAC <200). Each data point represents a gene. P-values were computed with the Wilcoxon non-parametric test.

### Supplementary Tables

| Supplementary Table 1. Summary statistics of the reference panels used in this study. | | | | | | | | |
|---|---|---|---|---|---|---|---|---|
| | **UKB WGS, autosomes** | **UKB WGS, chr20** | **HRC, chr20** | **KGP, chr20** | **Simulated 2.5k, 10Mb** | **Simulated 10k, 10Mb** | **Simulated 100k, 10Mb** | **Simulated 1M, 10Mb** |
| Number of samples | 140,119 | 140,119 | 27,165 | 2,504 | 2,500 | 10,000 | 100,000 | 1,000,000 |
| Number of variants | 582,534,516 | 13,296,637 | 864,277 | 2,140,589 | 96,737 | 223,116 | 747,162 | 2,274,530 |
| Number of rare < 0.1%MAF | 561,893,397 | 12,822,228 | 502,487 | 1,557,360 | 74,058 | 200,595 | 725,105 | 2,253,034 |
| Number of common ≥ 0.1%MAF | 20,641,119 | 474,409 | 361,790 | 583,229 | 22,679 | 22,521 | 22,057 | 21,496 |

| Supplementary Table 2. KGP reference panel. Imputation accuracy of chromosome 20 for 100 British samples. | | | | | | | |
|---|---|---|---|---|---|---|---|
| Ref. panel | Cov. | Chunk ID | Mean MAF | Number of SNPs | QUILT v1.0.4 | GLIMPSE v1.1.1 | GLIMPSE v2.0.0 |
| KGP | 0.1x | 0 | 0.00132952 | 244241 | 0.456408 | 0.411803 | 0.465187 |
| KGP | 0.1x | 1 | 0.00787683 | 66050 | 0.577527 | 0.51149 | 0.573788 |
| KGP | 0.1x | 2 | 0.041242 | 54644 | 0.745359 | 0.692588 | 0.734661 |
| KGP | 0.1x | 3 | 0.133536 | 42456 | 0.814958 | 0.776142 | 0.805226 |
| KGP | 0.1x | 4 | 0.293525 | 47418 | 0.836648 | 0.802387 | 0.826454 |
| KGP | 0.1x | 5 | 0.4487 | 21593 | 0.834323 | 0.801435 | 0.824797 |
| KGP | 1.0x | 0 | 0.00132952 | 244241 | 0.739091 | 0.724096 | 0.739482 |
| KGP | 1.0x | 1 | 0.00787683 | 66050 | 0.90473 | 0.896209 | 0.906514 |
| KGP | 1.0x | 2 | 0.041242 | 54644 | 0.966664 | 0.962374 | 0.965967 |
| KGP | 1.0x | 3 | 0.133536 | 42456 | 0.983718 | 0.981422 | 0.982613 |
| KGP | 1.0x | 4 | 0.293525 | 47418 | 0.986874 | 0.984594 | 0.985246 |
| KGP | 1.0x | 5 | 0.4487 | 21593 | 0.985793 | 0.983688 | 0.984296 |
| KGP | 4.0x | 0 | 0.00132952 | 244241 | 0.869276 | 0.856123 | 0.858291 |
| KGP | 4.0x | 1 | 0.00787683 | 66050 | 0.971135 | 0.970854 | 0.973916 |
| KGP | 4.0x | 2 | 0.041242 | 54644 | 0.990124 | 0.990422 | 0.99101 |
| KGP | 4.0x | 3 | 0.133536 | 42456 | 0.994693 | 0.99474 | 0.995015 |
| KGP | 4.0x | 4 | 0.293525 | 47418 | 0.995586 | 0.995537 | 0.99571 |
| KGP | 4.0x | 5 | 0.4487 | 21593 | 0.994728 | 0.994799 | 0.994877 |

| Supplementary Table 3. HRC reference panel. Imputation accuracy of chromosome 20 for 100 British samples | | | | | | | |
|---|---|---|---|---|---|---|---|
| Ref. panel | Cov. | Chunk ID | Mean MAF | Number of SNPs | QUILT v1.0.4 | GLIMPSE v1.1.1 | GLIMPSE v2.0.0 |
| HRC | 0.1x | 0 | 0.000120269 | 224227 | 0.207014 | 0.125994 | 0.199775 |
| HRC | 0.1x | 1 | 0.00027183 | 120818 | 0.351104 | 0.261677 | 0.343219 |
| HRC | 0.1x | 2 | 0.000602425 | 126696 | 0.481448 | 0.379143 | 0.478641 |
| HRC | 0.1x | 3 | 0.00132333 | 76873 | 0.565588 | 0.46726 | 0.560253 |
| HRC | 0.1x | 4 | 0.00262212 | 58452 | 0.609105 | 0.518484 | 0.605877 |
| HRC | 0.1x | 5 | 0.00576928 | 50362 | 0.669649 | 0.572071 | 0.666941 |
| HRC | 0.1x | 6 | 0.0130902 | 24282 | 0.700731 | 0.603178 | 0.695117 |
| HRC | 0.1x | 7 | 0.0264833 | 22831 | 0.779888 | 0.694648 | 0.77041 |
| HRC | 0.1x | 8 | 0.061651 | 32148 | 0.830007 | 0.760032 | 0.820082 |
| HRC | 0.1x | 9 | 0.13488 | 30553 | 0.857964 | 0.800864 | 0.850053 |
| HRC | 0.1x | 10 | 0.33348 | 68944 | 0.879336 | 0.833252 | 0.872496 |
| HRC | 1.0x | 0 | 0.000120269 | 224227 | 0.484692 | 0.457516 | 0.472836 |
| HRC | 1.0x | 1 | 0.00027183 | 120818 | 0.677793 | 0.653184 | 0.673778 |
| HRC | 1.0x | 2 | 0.000602425 | 126696 | 0.795029 | 0.769474 | 0.796396 |
| HRC | 1.0x | 3 | 0.00132333 | 76873 | 0.864126 | 0.843338 | 0.862137 |
| HRC | 1.0x | 4 | 0.00262212 | 58452 | 0.861404 | 0.841287 | 0.866068 |
| HRC | 1.0x | 5 | 0.00576928 | 50362 | 0.919274 | 0.902168 | 0.919566 |
| HRC | 1.0x | 6 | 0.0130902 | 24282 | 0.932108 | 0.916476 | 0.933445 |
| HRC | 1.0x | 7 | 0.0264833 | 22831 | 0.96623 | 0.957832 | 0.966741 |
| HRC | 1.0x | 8 | 0.061651 | 32148 | 0.978226 | 0.973191 | 0.978529 |
| HRC | 1.0x | 9 | 0.13488 | 30553 | 0.987454 | 0.984891 | 0.987211 |
| HRC | 1.0x | 10 | 0.33348 | 68944 | 0.990374 | 0.988452 | 0.990187 |
| HRC | 4.0x | 0 | 0.000120269 | 224227 | 0.656563 | 0.700222 | 0.687448 |
| HRC | 4.0x | 1 | 0.00027183 | 120818 | 0.813697 | 0.82628 | 0.829473 |
| HRC | 4.0x | 2 | 0.000602425 | 126696 | 0.895978 | 0.892986 | 0.90185 |
| HRC | 4.0x | 3 | 0.00132333 | 76873 | 0.931496 | 0.929783 | 0.939055 |
| HRC | 4.0x | 4 | 0.00262212 | 58452 | 0.915151 | 0.904321 | 0.927875 |

| | | | | | | | |
|---|---|---|---|---|---|---|---|
| HRC | 4.0x | 5 | 0.0057692 8 | 50362 | 0.965069 | 0.960919 | 0.968957 |
| HRC | 4.0x | 6 | 0.0130902 | 24282 | 0.966653 | 0.95928 | 0.970754 |
| HRC | 4.0x | 7 | 0.0264833 | 22831 | 0.98729 | 0.984486 | 0.987581 |
| HRC | 4.0x | 8 | 0.061651 | 32148 | 0.990441 | 0.987934 | 0.990796 |
| HRC | 4.0x | 9 | 0.13488 | 30553 | 0.994813 | 0.994368 | 0.994897 |
| HRC | 4.0x | 10 | 0.33348 | 68944 | 0.995913 | 0.995567 | 0.996024 |

| Supplementary Table 4. UKB reference panel. Imputation accuracy of chromosome 20 for 100 British samples | | | | | | | |
|---|---|---|---|---|---|---|---|
| Ref. panel | Cov. | Chunk ID | Mean MAF | Number of SNPs | QUILT v1.0.4 | GLIMPSE v1.1.1 | GLIMPSE v2.0.0 |
| UKB | 0.1x | 0 | 3.78E-06 | 3553687 | 0.475435 | 0.313571 | 0.457421 |
| UKB | 0.1x | 1 | 1.11E-05 | 2594376 | 0.509456 | 0.318188 | 0.502871 |
| UKB | 0.1x | 2 | 2.86E-05 | 734633 | 0.623176 | 0.433733 | 0.614108 |
| UKB | 0.1x | 3 | 5.50E-05 | 472654 | 0.68116 | 0.510873 | 0.674891 |
| UKB | 0.1x | 4 | 0.000118799 | 338275 | 0.727508 | 0.580628 | 0.725055 |
| UKB | 0.1x | 5 | 0.000265329 | 126256 | 0.779274 | 0.644206 | 0.778265 |
| UKB | 0.1x | 6 | 0.000529766 | 69910 | 0.76336 | 0.623869 | 0.7724 |
| UKB | 0.1x | 7 | 0.00119781 | 55926 | 0.73983 | 0.561406 | 0.749104 |
| UKB | 0.1x | 8 | 0.00270423 | 32105 | 0.756122 | 0.595768 | 0.778536 |
| UKB | 0.1x | 9 | 0.00540721 | 27074 | 0.783638 | 0.621338 | 0.794666 |
| UKB | 0.1x | 10 | 0.0122699 | 31024 | 0.816731 | 0.673628 | 0.817203 |
| UKB | 0.1x | 11 | 0.0273116 | 22484 | 0.862428 | 0.754799 | 0.865294 |
| UKB | 0.1x | 12 | 0.0550293 | 22873 | 0.889831 | 0.796489 | 0.887941 |
| UKB | 0.1x | 13 | 0.126886 | 39400 | 0.908759 | 0.835717 | 0.908556 |
| UKB | 0.1x | 14 | 0.278058 | 42977 | 0.918356 | 0.855346 | 0.919343 |
| UKB | 0.1x | 15 | 0.438644 | 24374 | 0.915326 | 0.851263 | 0.914515 |
| UKB | 1.0x | 0 | 3.78E-06 | 3553687 | 0.662739 | 0.360218 | 0.602821 |
| UKB | 1.0x | 1 | 1.11E-05 | 2594376 | 0.740284 | 0.577756 | 0.735726 |
| UKB | 1.0x | 2 | 2.86E-05 | 734633 | 0.846655 | 0.747809 | 0.848584 |
| UKB | 1.0x | 3 | 5.50E-05 | 472654 | 0.894341 | 0.82889 | 0.895097 |
| UKB | 1.0x | 4 | 0.000118799 | 338275 | 0.925349 | 0.89198 | 0.926687 |
| UKB | 1.0x | 5 | 0.000265329 | 126256 | 0.939118 | 0.92869 | 0.942045 |
| UKB | 1.0x | 6 | 0.000529766 | 69910 | 0.94575 | 0.933279 | 0.94828 |
| UKB | 1.0x | 7 | 0.00119781 | 55926 | 0.947631 | 0.945944 | 0.952721 |
| UKB | 1.0x | 8 | 0.00270423 | 32105 | 0.949099 | 0.957252 | 0.954474 |
| UKB | 1.0x | 9 | 0.00540721 | 27074 | 0.966497 | 0.969517 | 0.971006 |
| UKB | 1.0x | 10 | 0.0122699 | 31024 | 0.972889 | 0.969515 | 0.972233 |
| UKB | 1.0x | 11 | 0.0273116 | 22484 | 0.984621 | 0.984764 | 0.986229 |
| UKB | 1.0x | 12 | 0.0550293 | 22873 | 0.990007 | 0.990737 | 0.990886 |
| UKB | 1.0x | 13 | 0.126886 | 39400 | 0.993657 | 0.993912 | 0.993905 |

| | | | | | | | |
|---|---|---|---|---|---|---|---|
| UKB | 1.0x | 14 | 0.278058 | 42977 | 0.995469 | 0.99584 | 0.995667 |
| UKB | 1.0x | 15 | 0.438644 | 24374 | 0.994957 | 0.994966 | 0.994791 |
| UKB | 4.0x | 0 | 3.78E-06 | 3553687 | 0.788661 | 0.312655 | 0.689888 |
| UKB | 4.0x | 1 | 1.11E-05 | 2594376 | 0.818656 | 0.564381 | 0.812703 |
| UKB | 4.0x | 2 | 2.86E-05 | 734633 | 0.899196 | 0.69028 | 0.895555 |
| UKB | 4.0x | 3 | 5.50E-05 | 472654 | 0.931744 | 0.788517 | 0.931624 |
| UKB | 4.0x | 4 | 0.000118799 | 338275 | 0.952584 | 0.853959 | 0.948483 |
| UKB | 4.0x | 5 | 0.000265329 | 126256 | 0.962305 | 0.905157 | 0.954744 |
| UKB | 4.0x | 6 | 0.000529766 | 69910 | 0.966837 | 0.927129 | 0.960433 |
| UKB | 4.0x | 7 | 0.00119781 | 55926 | 0.967385 | 0.951563 | 0.961294 |
| UKB | 4.0x | 8 | 0.00270423 | 32105 | 0.958122 | 0.948099 | 0.950109 |
| UKB | 4.0x | 9 | 0.00540721 | 27074 | 0.977031 | 0.975357 | 0.973269 |
| UKB | 4.0x | 10 | 0.0122699 | 31024 | 0.983447 | 0.974791 | 0.977289 |
| UKB | 4.0x | 11 | 0.0273116 | 22484 | 0.990013 | 0.989221 | 0.989302 |
| UKB | 4.0x | 12 | 0.0550293 | 22873 | 0.992419 | 0.992703 | 0.992205 |

| Supplementary Table 5. KGP reference panel. Imputation running times and cost estimates of chromosome 20 for 100 British samples on the UKB RAP. Underlined cost represent the cost paid on the UKB RAP with normal priority. Bold lines represent the most efficient method. Estimates of whole genome costs per samples are based on the instances used in normal priority. | | | | | | | | |
|---|---|---|---|---|---|---|---|---|
| Software | Ref. panel | Cov. | Jobs | Virtual machine | Total time for chr20 (hh:mm:ss) | Cost on demand (GBP) | Cost on spot (GBP) | Whole genome cost per sample (GBP) |
| QUILT v1.0.4 | KGP | 0.1x | 32 | mem1_ssd1_v2_x4 | 10:15:35 | 1.02 | 0.27 | 0.14 |
| GLIMPSE v1.1.1 | KGP | 0.1x | 32 | mem1_ssd1_v2_x4 | 13:04:13 | 1.3 | 0.35 | 0.17 |
| **GLIMPSE v2.0.0** | **KGP** | **0.1x** | **16** | **mem1_ssd1_v2_x4** | **04:19:28** | **0.43** | **0.11** | **0.06** |
| QUILT v1.0.4 | KGP | 1.0x | 32 | mem1_ssd1_v2_x4 | 23:35:20 | 2.34 | 0.62 | 0.31 |
| GLIMPSE v1.1.1 | KGP | 1.0x | 32 | mem1_ssd1_v2_x4 | 11:26:05 | 1.13 | 0.3 | 0.15 |
| **GLIMPSE v2.0.0** | **KGP** | **1.0x** | **16** | **mem1_ssd1_v2_x4** | **06:01:10** | **0.6** | **0.16** | **0.08** |
| QUILT v1.0.4 | KGP | 4.0x | 32 | mem1_ssd1_v2_x4 | 58:35:13 | 5.81 | 1.55 | 0.77 |
| GLIMPSE v1.1.1 | KGP | 4.0x | 32 | mem1_ssd1_v2_x4 | 08:59:36 | 0.89 | 0.24 | 0.12 |
| **GLIMPSE v2.0.0** | **KGP** | **4.0x** | **16** | **mem1_ssd1_v2_x4** | **06:24:20** | **0.64** | **0.17** | **0.08** |

**Supplementary Table 6. HRC reference panel. Imputation running times and cost estimates of chromosome 20 for 100 British samples on the UKB RAP. Underlined cost represent the cost paid on the UKB RAP with normal priority. Bold lines represent the most efficient method. Estimates of whole genome costs per samples are based on the instances used in normal priority.**

| Software | Ref. panel | Cov. | Jobs | Virtual machine | Total time for chr20 (hh:mm:ss) | Cost on demand (GBP) | Cost on spot (GBP) | Whole genome cost per sample (GBP) |
|---|---|---|---|---|---|---|---|---|
| QUILT v1.0.4 | HRC | 0.1x | 32 | mem1_ssd1_v2_x4 | 12:05:25 | 1.2 | 0.32 | 0.16 |
| GLIMPSE v1.1.1 | HRC | 0.1x | 32 | mem1_ssd1_v2_x4 | 07:37:48 | 0.76 | 0.2 | 0.1 |
| **GLIMPSE v2.0.0** | **HRC** | **0.1x** | **16** | **mem1_ssd1_v2_x4** | **02:57:59** | **0.29** | **0.08** | **0.04** |
| QUILT v1.0.4 | HRC | 1.0x | 32 | mem1_ssd1_v2_x4 | 24:00:48 | 2.38 | 0.63 | 0.32 |
| GLIMPSE v1.1.1 | HRC | 1.0x | 32 | mem1_ssd1_v2_x4 | 04:36:50 | 0.46 | 0.12 | 0.06 |
| **GLIMPSE v2.0.0** | **HRC** | **1.0x** | **16** | **mem1_ssd1_v2_x4** | **03:52:58** | **0.39** | **0.1** | **0.05** |
| QUILT v1.0.4 | HRC | 4.0x | 32 | mem1_ssd1_v2_x4 | 49:53:50 | 4.95 | 1.32 | 0.66 |
| GLIMPSE v1.1.1 | **HRC** | **4.0x** | **32** | **mem1_ssd1_v2_x4** | **03:38:11** | **0.36** | **0.1** | **0.05** |
| **GLIMPSE v2.0.0** | HRC | 4.0x | 16 | mem1_ssd1_v2_x4 | 04:21:58 | 0.43 | 0.12 | 0.06 |

| Supplementary Table 7. UKB reference panel. Imputation running times and cost estimates of chromosome 20 for 100 British samples on the UKB RAP. Underlined cost represent the cost paid on the UKB RAP with normal priority. Bold lines represent the most efficient method. Estimates of whole genome costs per samples are based on the instances used in normal priority. | | | | | | | | |
|---|---|---|---|---|---|---|---|---|
| Software | Ref. panel | Cov. | Jobs | Virtual machine | Total time for chr20 (hh:mm:ss) | Cost on demand (GBP) | Cost on spot (GBP) | Whole genome cost per sample (GBP) |
| QUILT v1.0.4 | UKB | 0.1x | 32 | mem3_ssd1_v2_x48 | 247:18:56 | 288.86 | 52.23 | 144.43 |
| GLIMPSE v1.1.1 | UKB | 0.1x | 32 | mem3_ssd1_v2_x4 | 84:23:21 | 12.32 | 2.23 | 1.11 |
| **GLIMPSE v2.0.0** | **UKB** | **0.1x** | **16** | **mem2_ssd1_v2_x4** | **04:52:55** | **0.55** | **0.13** | **0.06** |
| QUILT v1.0.4 | UKB | 1.0x | 32 | mem3_ssd1_v2_x48 | 415:45:13 | 485.6 | 87.81 | 242.8 |
| GLIMPSE v1.1.1 | UKB | 1.0x | 32 | mem3_ssd1_v2_x4 | 83:38:20 | 12.21 | 2.21 | 1.1 |
| **GLIMPSE v2.0.0** | **UKB** | **1.0x** | **16** | **mem2_ssd1_v2_x4** | **06:07:29** | **0.69** | **0.16** | **0.08** |
| QUILT v1.0.4 | UKB | 4.0x | 32 | mem3_ssd1_v2_x48 | 561:06:31 | 655.38 | 118.51 | 327.69 |
| GLIMPSE v1.1.1 | UKB | 4.0x | 32 | mem3_ssd1_v2_x4 | 80:05:16 | 11.69 | 2.11 | 1.06 |
| **GLIMPSE v2.0.0** | **UKB** | **4.0x** | **16** | **mem2_ssd1_v2_x4** | **06:36:31** | **0.75** | **0.17** | **0.09** |
